# The neurodevelopmental transcriptome of the *Drosophila melanogaster* microcephaly gene *abnormal spindle* reveals a role for temporal transcription factors and the immune system in regulating brain size

**DOI:** 10.1101/2023.01.09.523369

**Authors:** Maria C. Mannino, Mercedes Bartels Cassidy, Steven Florez, Zeid Rusan, Shalini Chakraborty, Todd Schoborg

## Abstract

The coordination of cellular behaviors during neurodevelopment is critical for determining the form, function, and size of the central nervous system. Mutations in the vertebrate *Abnormal Spindle-Like, Microcephaly Associated (ASPM)* gene and its *Drosophila melanogaster* ortholog *abnormal spindle (asp)* lead to microcephaly, a reduction in overall brain size whose etiology remains poorly defined. Here we provide the neurodevelopmental transcriptional landscape for a *Drosophila* model for autosomal recessive primary microcephaly (MCPH) and extend our findings into the functional realm in an attempt to identify the key cellular mechanisms responsible for Asp-dependent brain growth and development. We identify multiple transcriptomic signatures, including new patterns of co-expressed genes in the developing CNS. Defects in optic lobe neurogenesis were detected in larval brains through downregulation of temporal transcription factors (tTFs) and Notch signaling targets, which correlated with a significant reduction in brain size and total cell numbers during the neurogenic window of development. We also found inflammation as a hallmark of *asp* MCPH brains, detectable throughout every stage of CNS development, which also contributes to the brain size phenotype. Finally, we show that apoptosis is not a primary driver of the *asp* MCPH phenotype, further highlighting an intrinsic Asp-dependent neurogenesis promotion mechanism that is independent of cell death. Collectively, our results suggest that the etiology of *asp* MCPH is complex and that a comprehensive view of the cellular basis of the disorder requires an understanding of how multiple pathway inputs collectively determine the microcephaly phenotype.

**AUTHOR SUMMARY:** Autosomal recessive primary microcephaly (MCPH) is a neurodevelopmental disorder characterized by a reduction in brain size, intellect, and life span. Over 30 genes have been found mutated in human MCPH patients, with *Abnormal Spindle-Like, Microcephaly Associated (ASPM)* being the most common. Although the clinical aspects of the disorder are well-characterized, the underlying cellular and molecular mechanisms are not. The fruit fly, *Drosophila melanogaster,* has an ortholog of the *ASPM* gene named *abnormal spindle (asp),* and mutations also give rise to fruit flies with small brains. It had previously been suggested that mitotic spindle defects were responsible for the *asp* MCPH phenotype, preventing neural stem cells from dividing properly to generate the necessary number of neurons and glia in the brain. However, genetic studies in flies showed that this wasn’t the case, suggesting that our knowledge of MCPH remains incomplete and must be revised. In this manuscript, we identified new pathways important for *asp-*dependent brain growth through transcriptional profiling. We found a number of key pathways disrupted in *asp* mutants, which have not been described previously. Using genetic tools in the fly, we tested a subset of these pathways to identify their contributions to the brain growth defects seen in *asp* mutants. Our results add to the growing number of pathways necessary for brain growth control, and provide a suitable foundation for follow-up genetic studies to assess MCPH.

## INTRODUCTION

Autosomal recessive primary microcephaly (MCPH) is a congenital neurodevelopmental disorder characterized by a reduction in overall brain size, intellectual disabilities, and shortened life span [1]. While the clinical manifestations of the disorder are well-characterized, the underlying molecular mechanisms responsible have been difficult to pinpoint. This is due to the increasing number of unique genes (∼30) that have been found mutated in human MCPH patients, each of which have known roles in diverse cellular functions such as mitosis, centriole structure and function, DNA damage and repair, chromatin remodeling, and nuclear envelope integrity [2]. This genetic diversity implies that multiple cellular pathways can give rise to the small brain phenotype, although our understanding of how this occurs remains limited.

*Abnormal Spindle-Like, Microcephaly-Associated (ASPM)* is the most commonly mutated gene found in human MCPH patients, accounting for more than 50% of all reported cases [3]. The *Drosophila melanogaster* ortholog, *abnormal spindle (asp)*, was identified before its human counterpart and was originally described as an essential mitosis gene, required for proper mitotic spindle morphology, mitotic progression, and chromosome segregation [4]. Cell biology, genetic, and biochemical studies by multiple labs have since provided a molecular explanation for how Asp functions as a ‘glue’ to maintain spindle pole and centrosome-pole cohesion during metaphase by associating with microtubule minus ends, as well as having additional roles in actin cytoskeleton regulation that may be important for proper tissue architecture [5–10].

The observed defects in mitosis provide an intuitive hypothesis for how mutations in *asp/ASPM* lead to reduced brain size, which can be broadly summarized into a defective neurogenesis program [11]. Neurogenesis is the developmental process by which neural stem cells produce a final pool of fully differentiated, post-mitotic neurons and glia in the central nervous system (CNS) [12]. In primates, brain size follows linear cell scaling rules with cell number being the primary driver [13]. Thus, mechanisms that promote efficient neurogenesis such as error-free mitosis is thought to have a significant influence on total neuron and glia numbers, and ultimately final brain size.

Given Asp/ASPM’s prominent role in mitosis, much of the earlier work attempting to link the cell biology to MCPH focused on cell division defects in the etiology of the disorder. In mice, ASPM was shown to regulate mitotic spindle orientation, preventing a premature switching of proliferative symmetric divisions in neuroepithelial cells to neurogenic, asymmetric divisions to ensure the appropriate amount of cortex neurons and glia [14]. A similar defect in spindle orientation was also found in human cultured cells lacking ASPM, along with defects in cytokinesis [15]. *Drosophila asp* has additional roles in spindle pole focusing and centrosome-pole attachment, which led to delayed anaphase onset, extended mitosis, and errors in chromosome segregation which were thought to be responsible for the small brain phenotype in flies [5,7,8].

However, disrupted mitotic spindle morphology alone is not sufficient to explain the small brain phenotype in flies, as only the N-terminal half of the Asp protein was sufficient to rescue MCPH phenotypes despite the fact that mitotic spindle morphology was still disrupted in these animals [5,8,16]. Furthermore, recent vertebrate studies have suggested that ASPM has additional, non-mitotic functions that may the primary driver of the small brain phenotype [17, 18], suggesting that our knowledge of the cellular basis of *asp/ASPM* microcephaly is incomplete and likely involves a combination of both mitotic and interphase roles that contribute to the small brain phenotype.

Next Generation Sequencing and other –omics-based approaches can provide insight into the spectrum of biological pathways contributing to MCPH [19], yet they have been underutilized in the MCPH field owing to limitations in model systems and the difficulty associated with acquiring viable samples from human patients [20–23]. *Drosophila* provides a unique model to explore these pathways in a temporal fashion, as the bulk of neurogenesis, neuronal remodeling, and maintenance of the final CNS development program occur at distinct developmental stages [24], thus allowing for identification of the relevant cellular pathways involved and also when in the neurodevelopmental pipeline they become important for brain size determination. Furthermore, the fly genetic tools available allow researchers to extend their analysis beyond discovery and into functional testing of hypotheses based on high throughput data to better understand how multiple inputs are coordinated to affect CNS growth and development.

Here we provide the transcriptome for *asp* mutant brains during the larval, pupal, and adult stages of neurodevelopment. Using co-expression and functional network analysis, we identify multiple biological pathways and gene regulatory modules enriched in the datasets. We found a number of temporal transcription factors that are critical for optic lobe neurogenesis and neuronal diversity that were downregulated in larval brains, along with Notch signaling targets. This correlated with a significant reduction in brain size and CNS cell number during the larval stage of development, providing a plausible mechanism to link neurogenic defects to the MCPH phenotype. We also identify inflammatory signatures during all developmental stages, suggesting immune system activation may be a hallmark of *asp* MCPH. Genetic analysis revealed that this partly contributes to the size phenotype as well. Other signatures included proteolysis, stress, metabolic, and actin cytoskeleton-related pathways that were enriched in at least two of the three stages. Surprisingly, we also did not find apoptosis to be a primary driver of *asp* MCPH, lending further support to a model in which *asp* promotes proper neurogenesis and cell number through cell death-independent mechanisms. Together, our results suggest that multiple inputs contribute to the *asp* brain phenotype, and provide a platform for future functional studies to explore the relative contributions of these pathways in the etiology of MCPH.

## RESULTS

### *asp* microcephaly affects the neurogenic phase of development and correlates with a reduction in total brain cell number

We previously reported that *asp* mutant adult brain volume is significantly reduced compared to wildtype animals, with the optic lobes showing the greatest reduction in size based on micro-CT (μ-CT) analysis [16]. In this study, we also wanted to extend our volume analysis to the larval and pupal stages to better understand the onset of the MCPH volume decrease phenotype and highlight the neurodevelopmental time course of the disorder. Neurogenesis occurs during the embryonic and larval stages, where symmetrically dividing neuroepithelial cells delaminate and form the asymmetrically dividing neuroblasts (NBs), which self-renew and generate ganglion mother cells (GMCs) that will later produce neurons and glia. Embryonic neurogenesis accounts for 10% of the neurons and glia that will contribute to the adult CNS, whereas 90% are generated during the larval period following a brief NB quiescence state [25]. Remodeling of the CNS occurs primarily during the pupal stage, and the end result is an adult brain consisting of ∼200,000 cells. This represents a 1,900% increase in cell number compared to the ∼10,000 cells found in the 1^st^ instar larval brain [24, 26] (Figure 1A).

**Figure 1.**
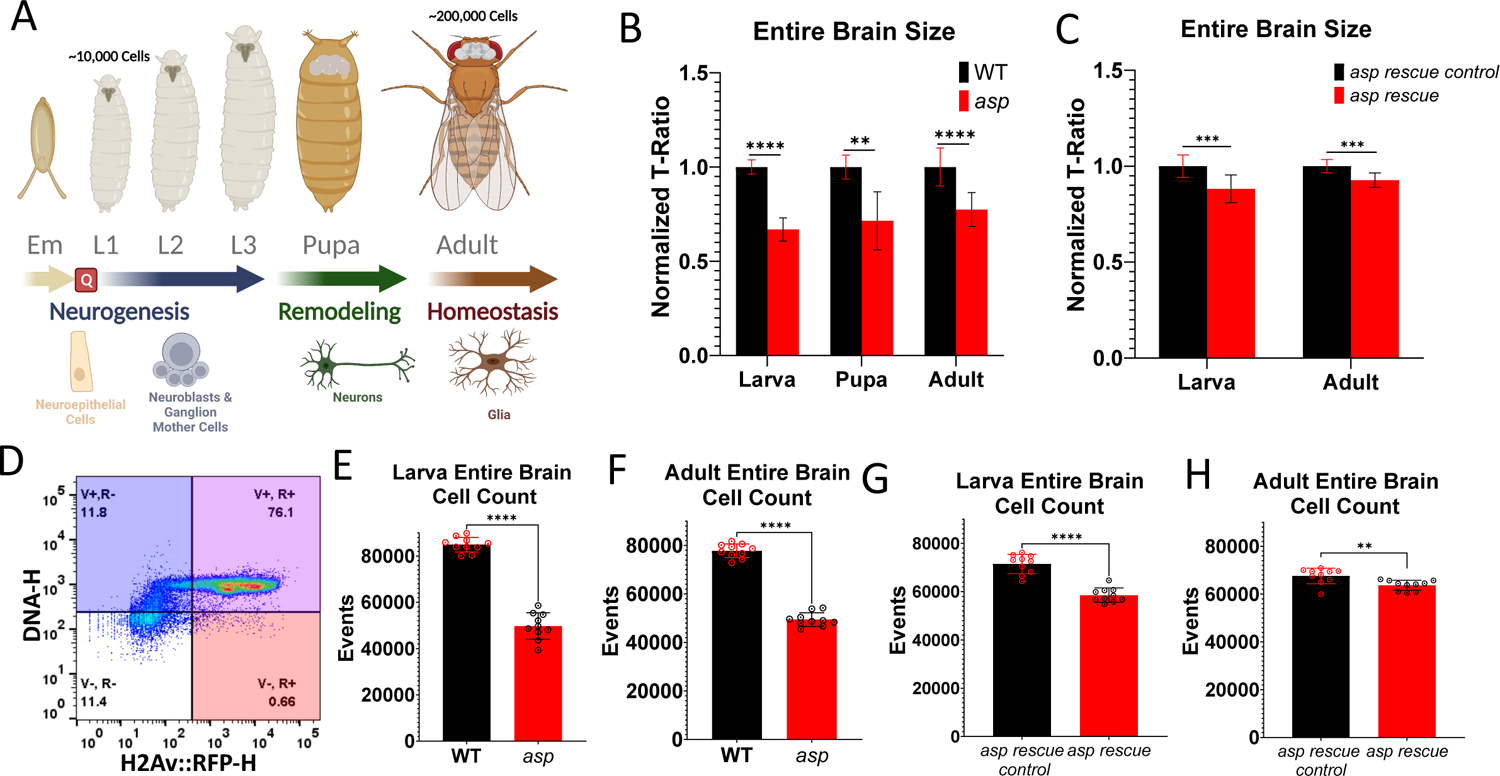
*asp* MCPH is a neurogenic disorder characterized by reduced brain size and total cell number. (A) Summary of fly neurodevelopment. Neurogenesis begins during the embryonic (Em) stage, where neuroepithelial cells delaminate from the neuroectoderm and form asymmetrically dividing neuroblasts. Neuroblasts divide in a self-renewing fashion to generate ganglion mother cells, which then further divide into post mitotic neurons and glia. A brief quiescent period (Q) precedes the larval neurogenesis period, with 90% of the cells present in the adult CNS generated during the L1, L2, and L3 stages. Neuronal remodeling occurs during the pupal stages, shaping the adult CNS into its final form consisting of ∼200,000 cells. (B) Microcomputed tomography (μ-CT) volume measurements from wildtype and *asp* mutant larva, pupa, and adults of the entire brain. (C) Volume measurements of the entire brain from *asp rescue* and *rescue control* animals at the larval and adult stages. (D) Validation of the flow cytometry method used to count neuronal cells from single brains. Animals expressing an RFP-tagged H2Av protein under control of the Act-Gal4 driver were co-labeled with DyeCycle Violet™. Events displaying high signal for both channels (V+, R+) were taken as the intact cell nuclei population and used to generate the counts displayed in (E) larval WT vs *asp mutant,* (F) adult WT vs *asp mutant,* (G) larval control and *asp rescue,* and (H) adult control and *asp rescue* brains. Each dot represents the count from a single brain (n=10). For all μ-CT measurements, data is represented as the T-ratio, which normalizes optic lobe volume to thorax width (body size) [16]. Wildtype (*asp^T25^/+*) was set to one and the subsequent genotypes were normalized accordingly. n≥5 brains, Welch’s t-test. ns, P>0.05; *P≤0.05; **P≤0.01; ***P≤0.001; ****P≤0.0001. Error bars represent standard deviation.

We chose three developmental time points to assess: 1) the late third instar larvae, when neural stem cells are actively dividing to generate neurons and glia (neurogenesis) [24]; 2) the pupal brain at ∼48 hrs APF when neuronal remodeling is occurring [27]; 3) the adult brain 3-5 days after eclosion when the majority of neurogenesis and remodeling have completed and thus reflects the end-point of neuronal development [28]. It is also equivalent to the developmental time point used to diagnose human MPCH patients in a clinical setting. A 33% and 28% decrease in the entire CNS in *asp* mutants compared to wildtype was observed in 3^rd^ instar larval and pupal brains, respectively (Figure 1B, Supplementary Figure 1A, 1B, 1C). We also found a 23% decrease in adult *asp* entire brain size that was highly significant (p<0.0001) (Figure 1B, Supplementary Figure 1E). A similar, but more pronounced trend was also observed when measuring optic lobe volume alone, with a 58%, and 42% reduction in size seen in pupal and adult brains, respectively (Supplementary Figure 1D, F). Given that the size decrease is readily evident towards the end of the neurogenic period (3^rd^ instar larva) and does not decrease further at later developmental stages, these results suggest that the *asp-*induced fly MCPH phenotype results primarily from neurogenic defects, analogous to its mammalian *ASPM* ortholog [14].

We also measured larval and adult brains from our previously described *asp* rescue strain [16]. The *asp* rescue strain expresses an N-terminally tagged GFP <u>m</u>inimal fragment (MF) of Asp consisting of the first 597 amino acids of the protein in the *asp* mutant background. Mitotic spindle morphology is still disrupted in *asp* rescue animals, with unfocused spindle poles and detached centrosomes evident, much like the *asp* mutants (Supplementary Figure 1J). However, this *asp* rescue fragment is able to restore both the entire brain and optic lobe sizes >90% in *asp* mutant adult brains, as well as neuropil morphology (Figure 1C, Supplementary Figure 1H, 1I) [16]. A similar trend (>88% rescue) was observed in the larval brain measurements as well (Figure 1C, Supplementary Figure 1G). These results suggests that the Asp^MF^ rescue fragment is largely capable of performing the necessary cellular roles of the full length protein that are required for proper brain size specification during the neurogenic window of development, which is independent of Asp’s role in mitotic spindle morphology.

To further investigate the cellular defects possibly contributing to the *asp* MCPH phenotypes, we performed flow cytometry on single fly brains (larval and adult) to determine total neuronal cell number. In primates, neuronal cell number is the primary driver of overall brain size [13]. Utilizing a fly brain-specific workflow outlined by the Buttitta lab [29], we validated its use in our hands and identified the relevant intact cell populations (Figure 1D, Supplementary Figure 2). We found a highly significant difference in cell number, with a 42% and 38% decrease in *asp* mutant third instar larva and adult brains, respectively, compared to wildtype (Figure 1E, 1F). We also measured neuronal cell number in *asp* rescue animals and found a similar trend to that observed in our μ-CT measurements of overall brain size: only an 18% and 6% decrease in cell number was observed in larval and adult *asp* rescue brains compared to *asp* rescue controls, respectively (Figure 1G, 1H). Together, these results confirm that *asp* mutant brains have reduced numbers of cells that can largely be restored by the *asp* rescue fragment, and that this reduction in cell number may be the primary driver of brain size in *Drosophila*, similar to primates [13].

### Transcriptional profiling of *asp* microcephaly brains throughout development

To identify genetic and cellular pathways that are disrupted in *asp* mutant brains and potentially contribute to the reduced number of cells and the microcephaly (MPCH) phenotype, we next analyzed the transcriptional profile of the central nervous system (CNS) from wildtype (*asp^T25^/+*), *asp* mutant (*asp^T25^/asp^Df^*), and *asp* rescue (*ubi-GFP::asp^MF^/+; asp^T25^/asp^Df^*) animals at the key developmental stages (third instar larva, P7 pupa, and 3-5 day old adults) used for the μ-CT and flow cytometry analysis above. Using this approach, we reasoned that we could identify the major genetic pathways contributing to MCPH and the key developmental stage when these defects occur to prevent proper brain growth (Figure 2A).

**Figure 2.**
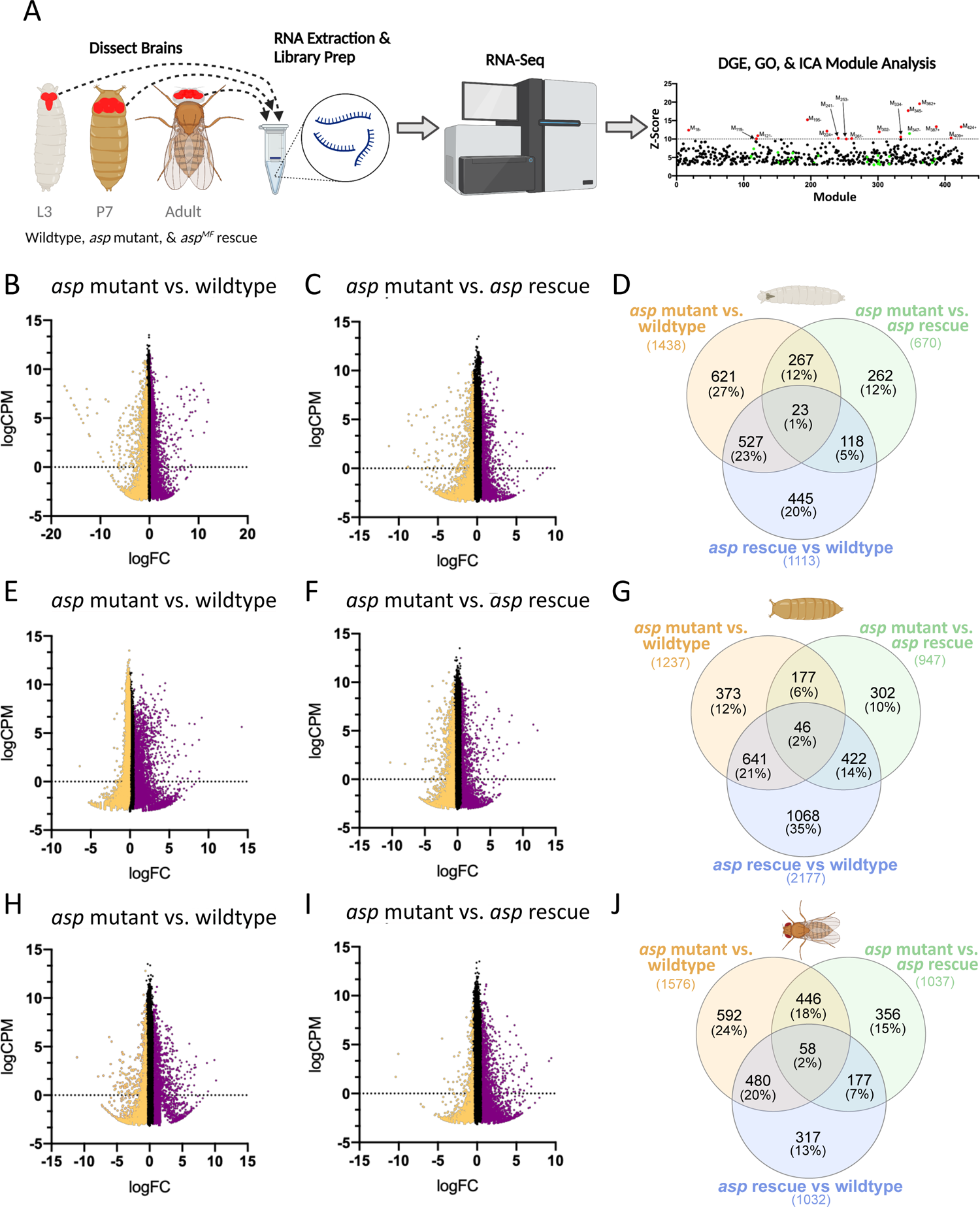
Identifying transcriptional signatures in a *Drosophila* model for microcephaly. (A) Experimental workflow for the brain RNA-Seq Analysis across multiple genotypes (WT, *asp* mutant, and *asp* rescue) and developmental stages (L3 wandering larvae, P7 pupae (∼48 hrs APF), and adults 3-5 days post eclosion). MA plots (logCPM vs logF.C.) for the *asp* mutant vs. wildtype DES analysis from (B) larvae, (E) pupae, and (H) adults. MA plots (logCPM vs logF.C.) for the *asp* mutant vs. *asp* rescue DES analysis from (C) larvae, (F) pupae, and (I) adults. Venn diagrams showing the percentage of shared and unique differentially expressed genes across *asp* mutant vs wildtype, *asp* mutant vs. *asp* rescue, and *asp* rescue vs. wildtype comparisons in (D) larvae, (G) pupae, and (J) adults.

After total RNA extraction, samples were subjected to Illumina sequencing and the resulting pair-end reads were aligned to the *D. melanogaster* genome. A total of 881 million pair-end reads were obtained; 663 million reads (78.1%) were mapped to the fly genome after adapter trimming and verifying sequence quality using FastQC. Biological replicates (n=4) clustered by genotype and developmental stage based on sample-to-sample distance and Principal Component Analysis (PCA), indicating suitable inter- and intra-group variation between the correct datasets for downstream analysis (Supplementary Figure 3).

### Impairment of neurogenic pathways and transcription factor networks in the *asp* mutant larval brain

We then performed pairwise differential gene expression (DGE) analyses for all genotype combinations at each developmental stage. Between ∼700 and 2,000 genes (coding and noncoding) were found to be significantly differentially expressed among the various combinations (logF.C. ≥0.5/≤-0.5; Adj. p-value ≤0.05) (Figure 2D, 2G, 2J, Supplementary File 1). Larval brains showed a roughly equal split between upregulated and downregulated genes across all genotype comparisons, while both pupal and adult brains showed a significant increase in upregulated genes compared to downregulated genes across *asp* mutant vs. wildtype and *asp* mutant vs. *asp* rescue comparisons (Figure 2B, 2C, 2E, 2F, 2H, 2I; Supplementary Figure 4). This suggests that the CNS transcriptome is dynamically regulated during development, and that *asp* mutations may significantly alter this landscape in a stage-specific manner.

To gain insight into the biological processes affected upon loss of *asp*, we utilized DAVID Functional Annotation to identify clusters of enriched Gene Ontology (GO) and KEGG pathway terms across multiple genotype comparisons and developmental stages (Supplementary File 2) [30, 31]. Our initial plan was to focus on the list of differentially expressed genes that were identified in both the *asp* mutant vs wildtype and *asp* mutant vs *asp* rescue analysis (Figure 2D, 2G, 2J). We reasoned that since brain size is largely restored in the *asp* rescue strain expressing the GFP::Asp minimal fragment (Asp^MF^) in the *asp* mutant background (Figure 1C) [16], it should act as a phenocopy for wildtype brains and thus the gene expression signature relevant to brain growth control between the two lines should be similar. This would allow us to eliminate any genes whose enrichment may have been due to genetic background differences (e.g., additional effects caused by the overexpression of the GFP::Asp minimal fragment) and enable identification of the conserved biological processes shared between both comparisons.

However, given the relatively low amount of gene overlap between these two genotype comparisons (267, 177, and 446 genes from larva, pupa, and adults, respectively (Figure 2D, 2G, 2J; Supplementary File 1), this analysis returned only a handful of statistically significant functional annotation clusters related to transcriptional regulation & transcription factor activity, stress responses, immune system, lipid metabolism, and cuticle development (Supplementary File 2). Therefore, we next applied DAVID Functional Annotation to the entire set of differentially expressed genes from the *asp* mutant (M) vs wildtype (W) (1,438 larval, 1,237 pupal, 1,576 adult genes) and the *asp* mutant (M) vs *asp* rescue (R) (670 larval, 947 pupal, 1,037 adult genes) comparisons (Figure 2D, 2G, 2J; Supplementary File 2). We then built an enrichment map network based on these functional clusters using EnrichmentMap in order to identify highly overlapping gene sets and networks between [M vs W] and [M vs R] at each developmental stage (Supplementary File 3) [32].

Several interesting networks were identified in this analysis. For larva, we noted a strong enrichment for transcriptional processes linked to neurogenesis, Notch signaling, and eye development and morphogenesis in the downregulated gene sets from [M vs W] and [M vs R] (Figure 3A). Shared genes enriched in the neurogenesis node included many known Notch players, including *Notch (N)* receptor, the downstream transcriptional targets belonging to the *enhancer of split (E(spl)* complex (e.g., *E(spl)mβ-HLH, E(spl)m4-BFM*) and the negative regulator *earmuff (erm)*. Other genes include the transcription factor *amos*, which promotes dendritic neuron formation; *H6-like homeobox (Hmx)*, which is involved in specification of neuronal cell types; and *off-track (otk)*, which is associated with non-canonical Wnt signaling (Supplementary File 3).

**Figure 3.**
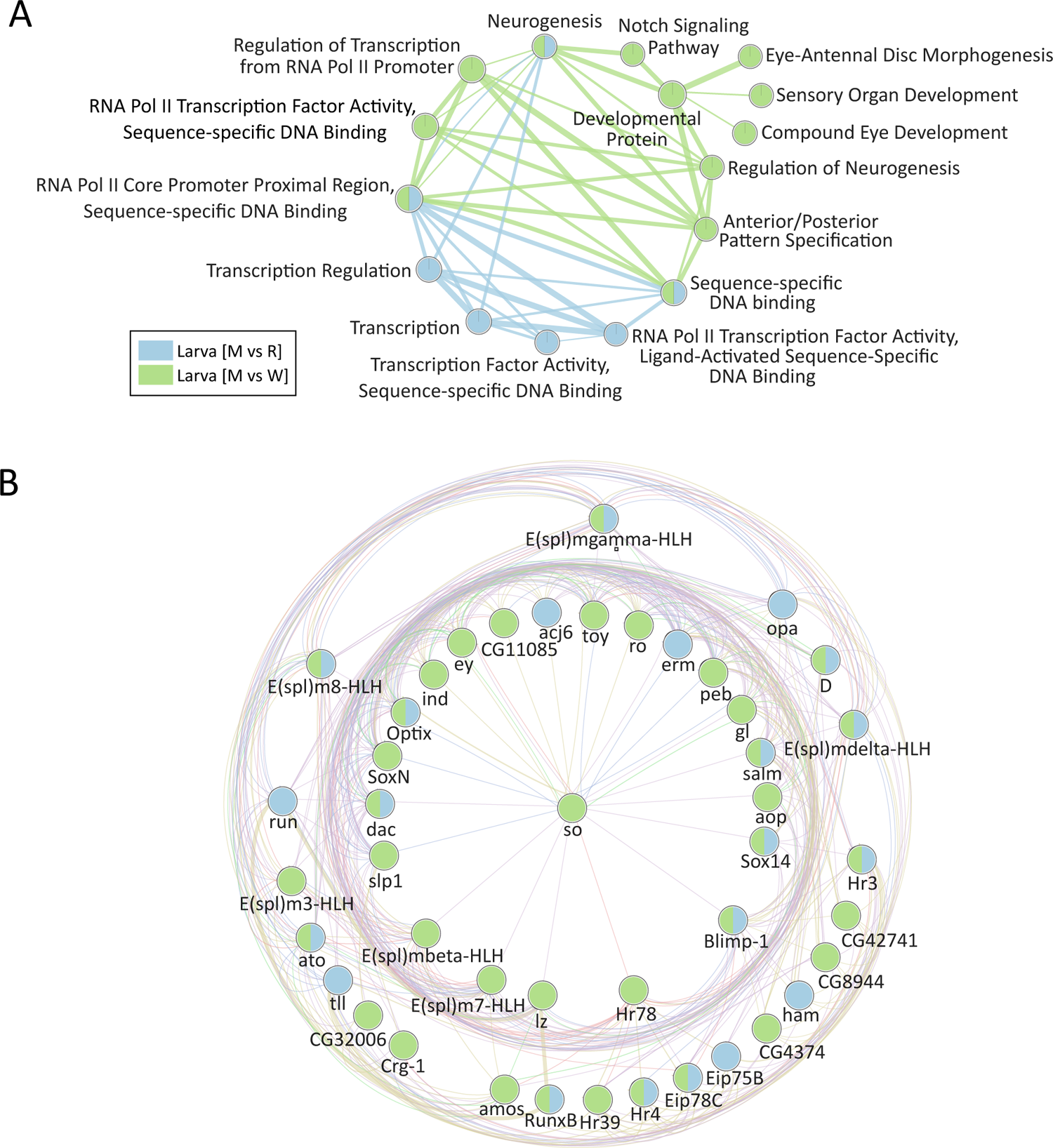
Transcription factor and signaling networks important for neurogenesis and tissue development are downregulated in *asp* mutant brains. EnrichmentMap visualization of the (A) transcription, neurogenesis, Notch signaling, and tissue development network identified from DAVID Functional Analysis of the *asp* mutant vs wildtype [M vs W] (green) and *asp* mutant vs *asp* rescue [M vs R] (blue) differentially expressed downregulated larval genes. Nodes represent a set of genes, edges (lines) represent gene overlap between each node. (B) Relationships between the 44 genes that comprise the “RNA Pol II Core Promoter Proximal Region, Sequence-specific DNA Binding” node as revealed by GeneMANIA. This gene set is highly enriched in spatial and temporal transcription factors that control optic lobe neurogenesis events. Line colors: green=genetic interactions; magenta=physical interactions; blue=co-localization; purple=co-expression; orange=predicted.

A strong enrichment for other well-known neurodevelopmental transcription factors involved in eye, optic lobe, and overall brain development was also observed in the larval comparison (Figure 3A, 3B), including a number of spatial and temporal TFs (tTFs) that are responsible for orchestrating the progression from neurogenesis to neuronal diversity in the fly visual system [33]. For example, the shared nodes labeled ‘RNA polymerase II core promoter proximal regional sequence-specific DNA binding’ (GO:0000978) and ‘Sequence specific DNA binding’ (GO:0043565) contain 44 and 28 genes, respectively. These genes include *Dichaete (D), tailless (tll), sloppy paired-1 (slp1), runt (run), eyeless (ey), odd paired (opa), Optix, twin of eyeless (toy), erm (earmuff), lozenge (loz), sine oculus (so), anterior open (aop), intermediate neuroblast defective (ind), hamlet (ham), glass (gla), and dachshund (dac)*, among many others, including the *E(Spl)* genes identified in the neurogenesis and Notch Signaling Pathway nodes (Figure 3B, Supplementary File 3). Only three of these transcription factors (*ato, Hr3, E(spl)mdelta*) were also found in the *asp* rescue vs wildtype comparison, suggesting that impaired neurogenesis through global downregulation of key neurodevelopmental spatial and temporal transcription factors and signaling pathways (e.g., Notch) may be a defining feature of *asp* mutant larval brains.

For the pupal and adult stages, only a few enriched downregulated networks were identified, in agreement with the finding that many fewer downregulated genes were available for this analysis (Supplementary Figure 4). Interestingly, transcription factor and chromatin related terms were also identified in the adult, including *Histone H4 replacement (His4r)*, *polybromo, early boundary activity 2 (Elba)*, and the notch-signaling related components *insensitive (insv) and transcription factor AP-2 (TfAP-2)*, while pupa had many fewer downregulated networks, primarily involving oxidoreductase activity (Supplementary File 3).

We next identified upregulated gene networks at each developmental stage between [M vs W] and [M vs R]. Glutathione and drug-related metabolic processes were the only shared networks observed in larval stages. However, several shared networks were found in the pupal and adult comparisons, including membrane, actin cytoskeleton, and chitin-based cuticle development networks. We also noted a shared, strong overlap for immune-related terms at these same developmental stages as well (Supplementary File 3).

### Inflammatory pathway activation may be a hallmark of *asp* mutant brains

While the gene network enrichments for the pairwise [M vs W] and [M vs R] comparisons at each developmental stage were useful for identifying stage-specific biological processes that may be contributing to *asp* MCPH, we next wanted to identify processes that were conserved across all developmental stages. This would allow us to determine if there is a common theme or hallmark of *asp* MCPH brains, which could then be tested genetically via mutational analysis to determine whether such shared processes are a cause or a consequence of the small brain phenotype. We therefore performed a similar gene network analysis ([M vs W] and [M vs R]) using EnrichmentMap, but did not separate these by developmental stage or expression (upregulated and downregulated). Thus, all six comparisons using the entire differentially expressed gene list were used for the analysis.

The results are shown in Figure 4. Nine networks were identified, including myosin and actin cytoskeleton, neuropeptide, glutathione & cytochrome P450 metabolism, chitin & cuticle development, proteolysis, sugar metabolism, membranes & cell junctions, oxidoreductases, and immune response (Figure 4A). However, the majority of these clusters consist of nodes that are only found in a few (≤3) of the pairwise comparisons and are therefore unlikely to serve as a hallmark of *asp* mutant brains. Thus, we focused on the networks that contained at least 4 of the 6 pairwise comparison for multiple nodes. This revealed a single cluster—immune response—as a potential hallmark of *asp* mutant brains throughout development (Figure 4A).

**Figure 4.**
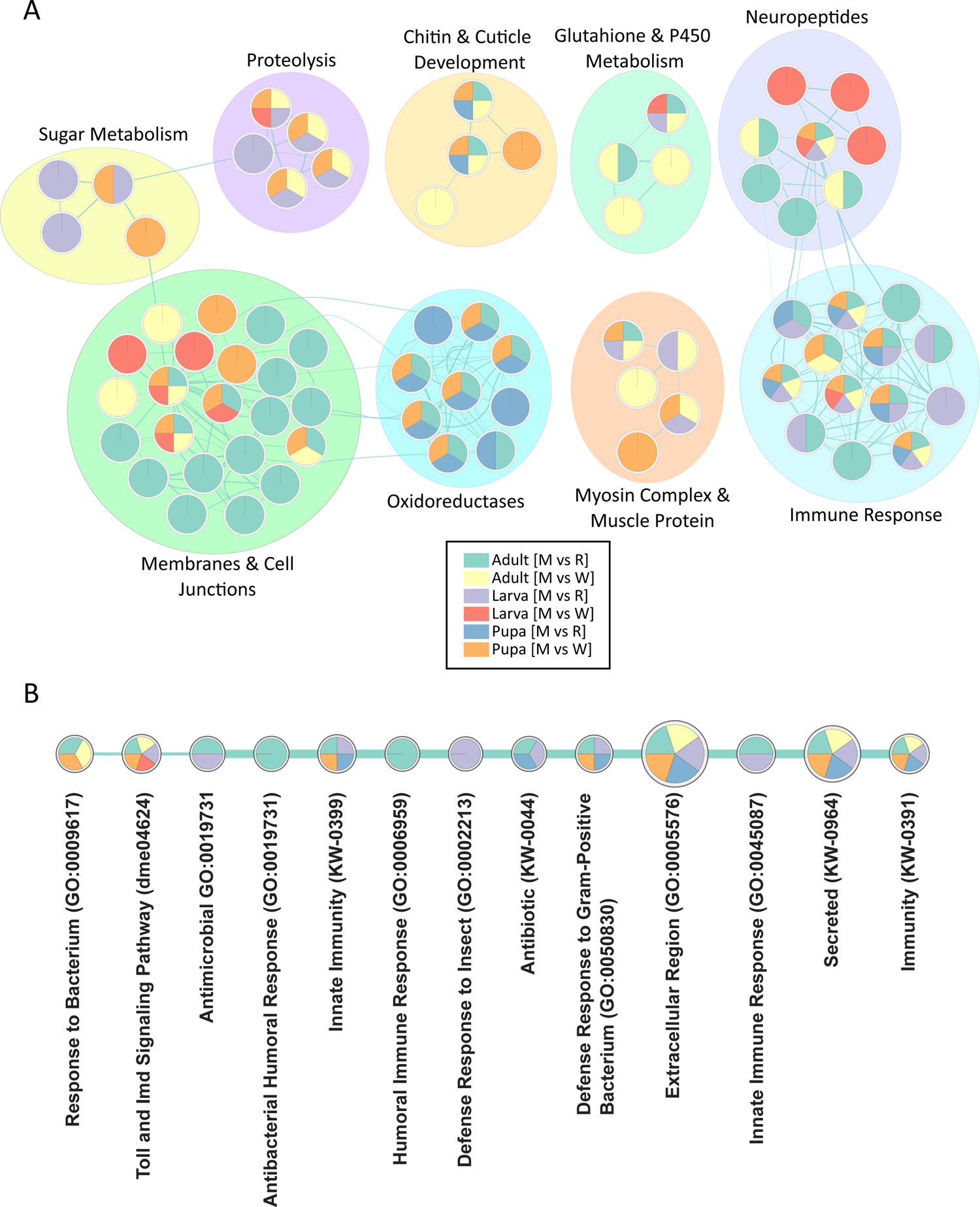
Biological networks conserved across developmental stages and genotype comparisons highlights inflammation as a hallmark of *asp* mutant brains. (A) EnrichmentMap clustering and visualization of nine networks identified as significantly enriched across larval, pupal, and adult brains from each genotype comparison. Node coloring for each genotype comparison is shown in the legend; nodes with multiple colors indicate the same DAVID Functional Annotation classification was found across multiple genotype comparisons and developmental stages. Individual node names are not shown for clarity (see Supplementary File 3 for a full list). (B) The immune response network visualized as a linear stack of nodes, with the name of each node representing the DAVID Functional Annotation identifier used to generate it (Gene Ontology (GO), KEGG pathway or Functional Annotation (KW)).

The nodes enriched in the immune system cluster are shown in Figure 4B, and consist of GO terms, KEGG pathways, and DAVID Functional Annotations that were used by EnrichmentMap to identify the Immune Response as a significantly enriched network. Node names such as ‘Toll and IMD Signaling Pathway’, ‘Innate Immunity’, ‘Defense Response to Gram-Positive Bacterium’, and ‘Immunity’ are especially prevalent across multiple genotype and developmental stage comparisons (Figure 4B). The ‘Immunity’ and ‘Toll and Imd Signaling Pathway’ gene sets contain the largest number of genes (44 and 38 genes, respectively) and are enriched in 5 out of the 6 pairwise comparisons. These genes include many well-known players in insect immunity, including the upstream receptors belonging to the *Peptidoglycan Recognition Protein (PGRP)* family, downstream effectors such as *pirk* and *IKKβ*, the activating NF-ΚB transcription factors *Relish (Rel), dorsal (dl),* and *Dorsal-related immunity factor (Dif),* and the downstream transcriptional targets belonging to the large family of secreted *Antimicrobial Peptides (AMPs)* (Cecropins, Diptericins, Attacins, Bomanins (IMs), etc.) (Supplementary File 3).

Further inspection of the immune-related nodes revealed two general trends. First, the immune response was found to be upregulated in virtually all of the pairwise comparisons, with the exception of the larval [M vs R] comparison. This had a significant enrichment of downregulated immune genes (∼10 total) belonging primarily to small subset of AMPs, such as *Attacin-A (AttcA)* and *Attacin-B (AttB)*. Secondly, we noted that the immune response appeared to increase as development proceeded, with a stronger and more consistent (e.g., upregulation) trend readily apparent in pupal and adult stages. These data suggests that an inflammatory response may be a hallmark of *asp*-induced microcephaly.

### Identifying enriched networks of co-regulated gene modules in *asp* mutant brains

To complement our functional enrichment analysis, we also utilized a *Drosophila*-specific gene signature set consisting of 850 modules of co-regulated genes identified through Independent Component Analysis (ICA) [34]. The advantage of this analysis is that it utilizes gene co-expression patterns identified in flies from thousands of high throughput datasets, and therefore may have additional analytical power to uncover subtle expression perturbations and the underlying gene regulatory networks. This is particularly true in the brain, where it has been used to identify transcriptional signatures unique to individual brain cell types [34] and thus can provide deeper insight into fly-specific brain function and physiology [19]. We therefore searched for enriched co-regulatory signatures that could be biologically relevant for MCPH.

The main strength of the ICA module analysis is that all gene logF.C.’s from a genotype pair are used when interrogating ICA modules rather than employing traditional cutoff values (based on logF.C. and adjusted p-value) as in DGE-based analyses. This maximizes expression pattern matching sensitivity for low amplitude (but nonetheless biologically relevant) signatures. In other words, small or subtle changes in logF.C. can be identified as significant if multiple genes part of a co-expressed cluster also have subtle changes and the majority move in the same direction (up or down regulated). This is illustrated by the M_393+_ module (Supplementary Figure 5B), which consists of over 400 co-expressed genes. 70% of these genes have logF.C.’s that do not meet the logF.C. ≥0.5/≤-0.5 criteria in both the [M vs W] and [M vs R] comparisons, yet the genes are given higher weights in this analysis and their module achieves a significant Z-score that meets the Z ≥ 3/≤-3 cutoff (Z =-4.9 and −4.4 for [M vs W] and [M vs R], respectively) as a result of their collective downregulation. Conversely, highly significant modules (Z-score ≥10/≤-10) consist of co-expressed genes with larger logF.C.’s (logF.C. ≥2/≤-2). This is illustrated by M_146-_, the most highly enriched shared module from the larval analysis. The majority of these genes have a large positive logF.C., and thus the module is assigned a highly significant positive Z-score (Z=15.4 and 18.3 for [M vs W] and [M vs R], respectively) (Figure 5A, Supplementary Figure 5A).

**Figure 5.**
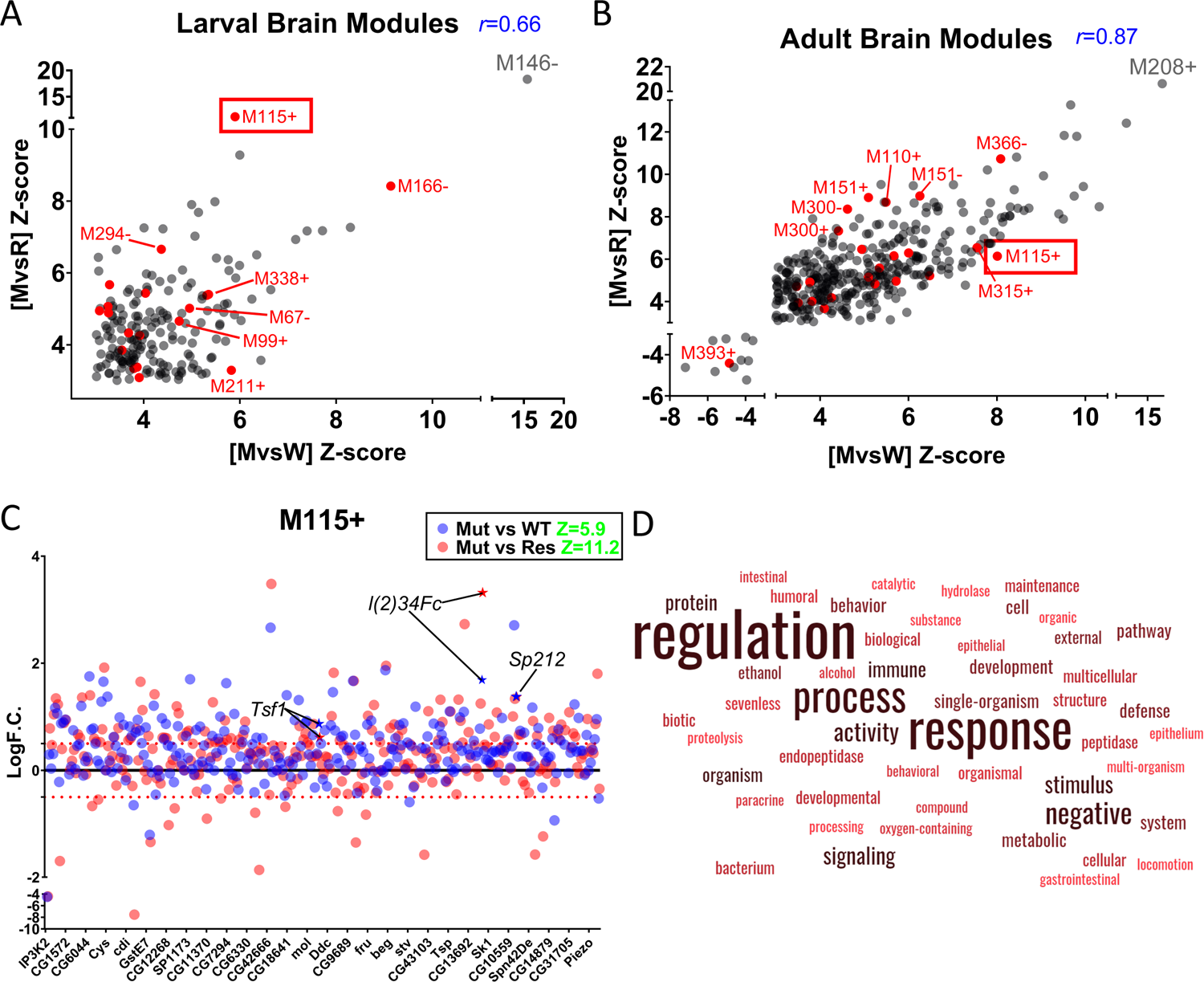
Identifying co-regulated gene network signatures in *asp* mutant brains. Scatterplots showing significantly enriched co-expression modules identified through ICA analysis of the [M vs W] and [M vs R] datasets from (A) larval and (B) adult brains. Each module is plotted as a point based on its Z-score enrichment from each analysis. Only significant modules with a Z-score of >3/<-3 are shown. Co-expressed modules containing at least one immune system-related GO term are colored red, with M_115+_ outlined with a red box. A subset of the more significantly enriched (Z-score ≥10) modules are also labeled in gray font. The Pearson’s Correlation Coefficient (*r)* is shown in blue font (two-tailed P<0.0001). (C) Plot of logF.C. vs. gene for the M_115+_ module. Z-score enrichment is shown in green font for each comparison. Each dot is colored based on the mutant vs wildtype (Mut vs WT, blue) and mutant vs rescue (Mut vs Res, red) value. Red dotted line indicates the 0.5/-0.5 logF.C. position. Not all gene names are included on the x-axis for clarity. Relevant immune system genes (*l(2)34Fc, Sp212, Tsf1*) are highlighted. (D) Word cloud enrichment of the GO terms associated with the M_115+_ co-expressed module, highly enriched for immune-related, development, and signaling words.

The summary of this analysis is shown in Figure 5 and Supplementary Figure 5, along with a file displaying the statistical enrichments and a description of the co-expressed genes and GO terms for each ICA module (Supplementary File 4). Using a Z-score cut-off of Z ≥ 3/≤-3, we evaluated both the [M vs W] and [M vs R] profiles individually as well as the intersection to find common co-expression patterns and filter out potential false positives. A total of 192, 98, and 377 modules met this criteria in larval, pupal, and adult stages, respectively. We also found a strong correlation between the significant modules identified in the [M vs W] and [M vs R] analyses and their Z-score magnitude (Pearson’s *r*= 0.66, 0.70, 0.87, p<0.0001 for larva, pupa and adult, respectively) (Figure 5A, 5B, Supplementary Figure 5D), further evidence that the *asp* rescue animals largely recapitulate the wildtype genotype. Interestingly, 97% of adult and 100% of larval and pupal modules were assigned positive Z-scores, meaning that the majority of the genes in each module were upregulated in the comparisons and is in general agreement with the overall expression patterns noted in each developmental stage (Figure 5A, 5B; Supplementary Figures 4A, 4B, 5D).

From an individual GO perspective, we found significant overlap with our EnrichmentMap Analysis (Figure 4A), thus validating both approaches. For example, M_146-,_ M_302-_ and M_208+_ were the most highly enriched (Z-score ≥8) shared modules in larva, pupa, and adult stages (Figure 5A, 5B; Supplementary Figure 5A, 5D) and contain genes involved in glutathione & sugar metabolism, transport, proteolysis, and cuticle development (Supplementary File 4). GO terms related to ‘neurogenesis’ were found in seven and twenty four modules enriched in larval [M vs W] and [M vs R] profiles, respectively, although only one of these neurogenesis modules were shared between both comparisons. This is also in agreement with our EnrichmentMap analysis (Figure 3).

Also, we again noted a very strong enrichment of modules containing GO terms related to the immune system at each developmental stage (red dots, Figure 5A, 5B, Supplementary Figure 5D). All of these modules, with the exception of adult M_393+,_ were assigned positive Z-scores in agreement with our earlier DGE findings that immune system genes are upregulated in the *asp* mutant. However, upon closer inspection of the immune genes found in the negatively-assigned M_393+_ cluster (27 total), there is a clear pattern of behavior where positive regulators of the immune response (e.g., *darkener of apricot (doa)* [35]) display positive logF.C.’s., and negative regulators (*Enhancer of bithorax (E(bx)), fat facets (faf)* [36, 37]) have negative logF.C.’s, consistent with an activated immune response in *asp* mutant adult brains (Supplementary Figure 5B). Furthermore, when we examined the 23 intersection modules that were conserved across larval, pupal, and adult brains, three (13%) of these were found to have a significant enrichment of co-expressed immune system genes and associated GO terms, with M_211+_ & M_99+_ enriched in many upregulated AMPs of the immune response (Supplementary File 4).

However, most biological outcomes require input from multiple genes, many of which have distinct functions (and therefore GO terms) that collectively contribute to the final output. This is the primary advantage of co-expression analysis, which is illustrated by one of the most highly enriched modules (M_115+_) shared across larval, pupal, and adult comparisons (Figure 5C). M_115+_ consists of a large number of co-expressed genes (>275), whose GO terms are enriched for words related to immunity (note the positive logF.C. values for the immune system genes *l(2)34Fc, Sp212,* and *Tsf1*), development, signaling, metabolism, proteolysis, etc., all of which were identified as separate networks in our EnrichmentMap analysis (Figure 4A, 5C, 5D). Although the biological significance of this co-expressed cluster requires further investigation, it suggests that not only are multiple genetic inputs with diverse functions required for *asp-* dependent brain growth control, but also that these processes may be coordinated at the transcriptional level to achieve the intended biological outcome in the CNS.

A second advantage of the co-expression analysis is identifying new patterns of transcription and potentially new roles for otherwise well-characterized genes. For example, the notch-responsive *e(spl)* genes that were highlighted in our EnrichmentMap analysis from the larval brain (Figure 3) were found in twelve different modules from the larval [M vs W] and [M vs R] intersection (M_166-_, M_202+_, M_164-_, M_148+_, M_186+_, M_178+_, M_113-_, M_397+_, M_399+_, M_15-_, M_107+_, M_140+_). Interestingly, M_166-_ was the second highest scoring module for the larva intersection (Z ≥ 8 for both) and contains only the notch-responsive *E(spl)mBeta-HLH* gene but no other *e(spl)* member (Figure 5A, Supplementary File 4). The other 217 genes in this module are involved in a diverse range of biological processes, including cell adhesion, actin cytoskeleton organization, immune system, ribonucleotide & nucleotide metabolism, and cell fate determination. However, 33% of the *e(spl)*-containing modules have no enriched GO-associated terms, because the majority of the genes present have no defined function (CG-designation on FlyBase). The best example of this is the *e(spl)mγ-HLH-*containing M_113-_ module, which contains 125 genes total, 70 of which are CG-genes (Supplementary Figure 5C, Supplementary File 4). This module is also interesting because it has positive Z-scores for both [M vs W] and [M vs R], driven by the majority of the CG-genes being upregulated in each comparison, yet *e(spl)mγ-HLH* is significantly downregulated (logF.C.’s of −1 and −1.5 in [M vs W] and [M vs R]) (Supplementary Figure 5C). Together, these results suggest that Notch signaling inputs may be coordinated with many other cellular pathways in parallel, whose identity and biological consequence remain open for investigation.

### Inflammation is detectable in *asp* brains but does not contribute to the tissue disorganization phenotype

Although our analyses highlighted multiple cellular pathways that are disrupted in *asp* mutant brains, the consistent identification of a transcriptional CNS immune response across development stages using multiple bioinformatics methods led us to further investigate inflammation’s role in *asp* MCPH. The canonical immune response in flies is mediated through two pathways, Toll and IMD, that mediate responses to various environmental triggers such as PAMPs (Pathogen-Associated Molecular Patterns (PAMPs) and DAMPs (Damage Associated Molecular Patterns) (Supplementary Figure 6A) [38]. Downstream activation of the Antimicrobial Peptides (AMPs) and other effectors occurs through a family of NF-ΚB transcription factors known as *relish (rel), dorsal (dl)*, and *dorsal-related immunity factor (Dif)* that mediate IMD (Rel) and Toll (dl and Dif) pathway activation [39].

Previous studies have linked immune system activation to neurodegenerative phenotypes seen in fly models of human neurodegeneration, such as ataxia-telangiectasia (A-T) [40, 41]. These phenotypes consist of vacuole-like structures (holes) in the brain that form as a result of hyper localized neuron and glial cell death and loss of the accompanying neuropil [42]. Hyper activation of the immune response through AMP production has been shown to induce death of dopaminergic neurons upon loss of *Cdk5*, and forced overexpression of AMPs independent of Toll or IMD pathway activation can also lead to extensive vacuole formation [43, 44]. Interestingly, we also observed vacuolar structures, prominently disrupted neuropil boundaries, and missing neuronal cell bodies in the optic lobe from *asp* mutant pupal and adult brains (Figure 6C, 6D, Supplementary Figure 6D, 6E). This led us to further test whether these phenotypes might be a direct result of immune system activation, and if so, determine their contribution to the MCPH phenotype.

**Figure 6.**
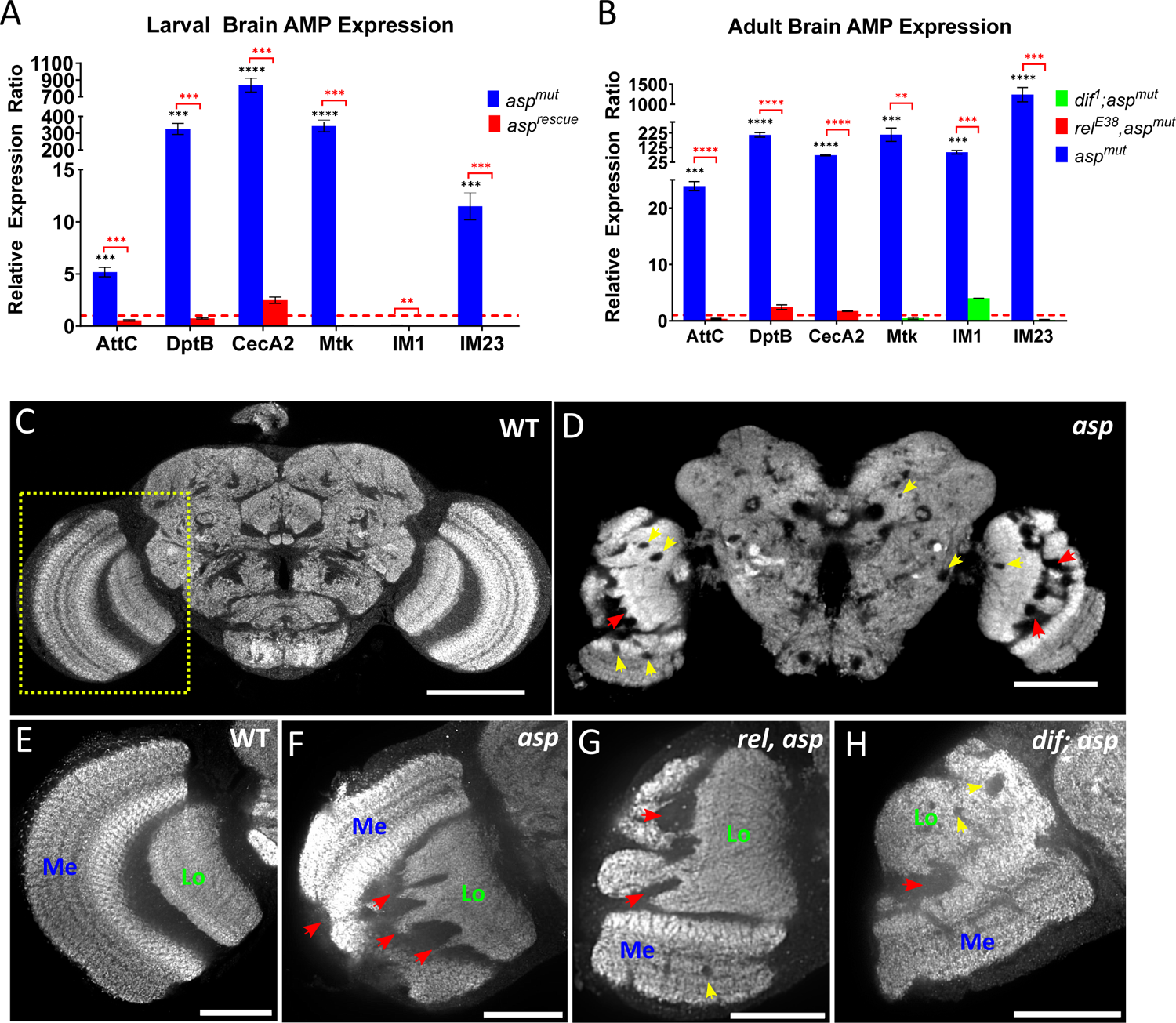
Immune system activation in *asp* mutant brains leads to upregulated AMP expression. (A) qPCR analysis of a subset of antimicrobial peptides (AMPs) identified as differentially expressed in the RNA-Seq analysis from larval brains. Data is shown as the relative expression ratio (RER), red dotted line indicates the RER value for wildtype, normalized to 1. Expression in *asp* mutant brains is shown in blue, *asp rescue* in red. (B) qPCR analysis in adults from the *asp* mutant (blue), *dif^1^; asp^mut^* (green), and *rel^E38^, asp^mut^* (red) genotypes showing suppression of AMP expression upon loss of NF-ΚB activity. Confocal imaging of adult brains labeled with α-brp (nc82) to visualize the neuropil in (C) wildtype and (D) *asp* mutants. Red arrowheads point out disrupted neuropil boundaries; yellow arrowheads indicated vacuole-like hole structures. The yellow dotted box indicates the adult optic lobe, which is shown in the insets from (E) wildtype, (F) *asp* mutant, (G) *rel^E38^, asp* double mutants, (H) *dif^1^; asp* double mutants. Medulla (Me) and Lobula (Lo) neuropil regions are labeled. ns, P>0.05; **P≤0.01; ***P≤0.001; ****P≤0.0001. Error bars represent standard deviation. Scale bars = 50 μm (E, G, H) and 100 μm (F).

We first verified that a subset of IMD and Toll pathway components were significantly upregulated in *asp* mutants using qPCR, in agreement with our RNA-Seq results (∼70% validation rate). Upstream components (*IMD, toll, rel, dif*) were mildly upregulated at different developmental stages (Supplementary Figure 6C). However, we observed significant upregulation of effector AMPs during both larval and adult stages (Figure 6A, 6B). IMD-regulated AMPs (*AttC, DptB, CecA2*) were consistently upregulated during both stages, with *CecA2* and *DptB* showing their largest upregulation during the larval period. Toll-regulated AMPs (*Mtk, IM1, IM23*) showed the strongest upregulation during the adult stage, with *IM23* expression increasing nearly 100x from larval to adult brains. AMP activation was also not observed in *asp* rescue animals, suggesting that this response was specific to *asp* mutants and not the result of Asp transgene overexpression (Figure 6A). Interestingly, we did not observe significant expression for these same AMPs during the pupal stage (Supplementary Figure 6B), suggesting that the AMP response is dynamically regulated throughout development in response to *asp* loss.

To test whether inflammation is a cause or consequence of *asp*-induced MCPH, we next generated double and triple mutant combinations of *asp* and the NF-ΚB factors *rel* (*rel^E38^*) and *dif (dif^1^)*. These NF-ΚB mutations suppressed AMP activation in a target-specific manner (Figure 6B), further validating the immune response seen in *asp* mutants. We first focused on the *asp* adult MCPH tissue morphology defects (abnormal neuropil projections, mis-positioned medulla, missing cell bodies, and vacuole formation) (Figure 6C, 6D), hypothesizing that if these defects were dependent on upregulated AMPs then their suppression in the NF-ΚB mutants should result in at least partial restoration of wildtype brain tissue. Qualitative evaluation of adult *rel, asp* and *dif, asp* double mutants compared to the single *asp* mutant did not reveal any significant tissue differences, with all mutant genotypes showing significant optic lobe neuropil disorganization and vacuole-like structures (Figure 6E-6H). Triple mutants (*dif; rel, asp*) also did not suppress the morphology defects of the *asp* single mutant (Supplementary Figure 6F, 6G). These results indicate the tissue morphology defects observed in *asp* microcephaly are not caused by AMP upregulation or other NF-ΚB-dependent gene regulatory networks.

### Genetic suppression of inflammation partially rescues the *asp* MCPH size phenotype

We next considered whether inflammation might contribute to the *asp* microcephaly brain size phenotype. Analysis of *rel, asp* and *dif, asp* double mutant combinations revealed a partial rescue of both entire brain and optic lobe volume compared directly to the *asp* single mutant alone (Figure 7A, 7B, Supplementary Figure 7A, 7B), with *rel, asp* and *dif; asp* double mutants showing a 28% and 26% increase in optic lobe volume, respectively (Figure 7B, Supplementary Figure 7B). This partial size rescue becomes even more apparent when factoring in the genetic background influences from the *rel* and *dif* single mutants. For example, only a 19% and 34% decrease in optic lobe volume was observed in the *rel, asp* and *dif; asp* double mutants compared to the *rel* and *dif* single mutants, respectively (note the 42% reduction seen in the *asp* single mutant compared to wildtype, Supplementary Figure 1F). The *dif; rel, asp* triple mutants showed a similar trend to each double mutant genotype, with a 29% increase in volume compared to the *asp* mutant and only a 27% reduction compared to the *dif* mutant (Figure 7A, 7B, Supplementary Figure 7A, 7B).

**Figure 7.**
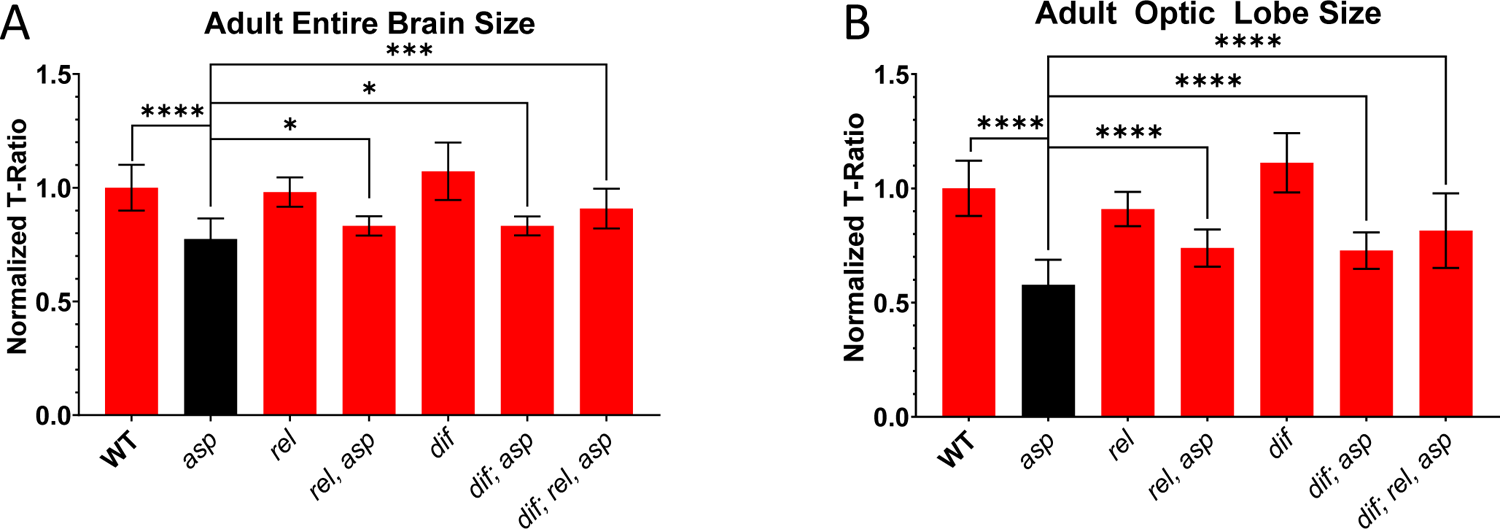
The NF-ΚB immunity factors *relish* and *dif* contribute to the *asp* microcephaly phenotype. Microcomputed tomography (μ-CT) volume measurements of (A) Entire brain and (B) optic lobe from WT, *asp* mutant, *asp, rel* double mutant*, asp, dif* double mutant, and *asp, rel, dif* triple mutants. For all graphs, data is represented as the T-ratio, which normalizes optic lobe volume to thorax width (body size) [16]. Wildtype (*asp^T25^/+*) was set to one and the subsequent genotypes were normalized accordingly. n≥5 brains, Welch’s t-test. ns, P>0.05; *P≤0.05; **P≤0.01; ***P≤0.001; ****P≤0.0001. Error bars represent standard deviation.

To understand the mechanism behind this partial suppression of the *asp* MCPH phenotype upon immune system inactivation, we focused on the role of glial cells, the primary modulator of the CNS immune response in flies and vertebrates [45]. *WDR62*, another microcephaly gene with roles during mitosis, was also shown to have a glial cell-specific function that is critical for promoting proper adult fly brain growth [46, 47]. We tested whether *asp* has a cell-autonomous function in glial cells by expression of either the *asp* rescue fragment (asp^MF^::GFP) or a full length copy (asp^FL^::GFP) with the pan-glial cell Gal4 driver *Repo.* We did not observe brain size rescue in these animals (Supplementary Figure 7C), suggesting Asp’s ability to promote proper brain size is not dependent on a glial cell-specific function, and its ability to potentially trigger inflammation in glial cells when mutated likely occurs non-cell autonomously.

Finally, we evaluated apoptosis, which can occur as a result of AMP activation in glial cells [40,41,43,48]. Cleaved Death Caspase-1 (DCP-1) staining did not reveal a significant difference in the total number of foci between *asp* mutant and wildtype larval brains, but we did find a ∼29% increase when normalizing to overall optic lobe volume, although this difference was not statistically significant (Figure 8A, 8B). Furthermore, genetically inhibiting apoptosis in *asp* mutants using the baculovirus anti-apoptotic protein P35 using either a neuroblast (*Insc-Gal4)* or neuron-specific (*Elav-Gal4*) Gal4 driver did not rescue either the brain size or morphology defects (Figure 8C, 8D, 8E). In fact, we detected a further significant 15% and 18% decrease in entire brain and optic lobe size in Elav-Gal4/UAS-P35 *asp* mutant animals compared to the *asp* mutant alone (Figure 8C, 8D). These results suggest that apoptosis is not the primary driver of the *asp* mutant MCPH phenotype, and that the downstream consequences of immune system activation in *asp* mutants that influence brain size remains unknown.

**Figure 8.**
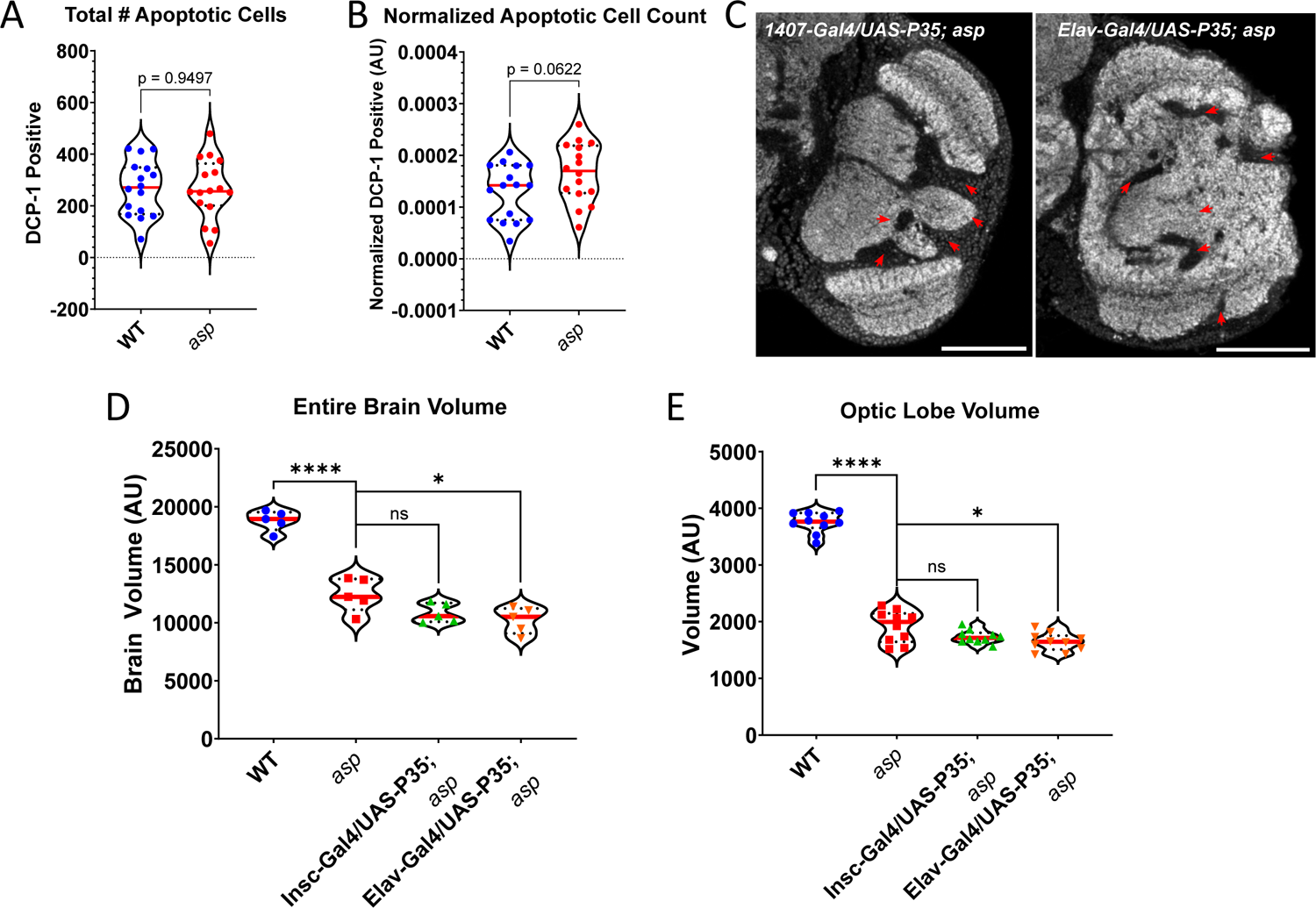
Apoptosis is elevated in *asp* mutants but does not contribute to the MCPH phenotype. (A) Total number of cleaved death caspase-1 (Dcp-1) positive foci in WT and *asp* mutant third instar larval brains. (B) Same analysis in (A), but normalized for optic lobe volume. (C) Adult brains stained with nc82 (α-brp) to visualize the neuropil of the optic lobe from *asp* mutants expressing the anti-apoptotic baculovirus UAS-P35 protein using the pan-neuronal Insc-Gal4 and neuron Elav-Gal4 driver. Note the severe morphological defects (red arrowheads) present, identical to those seen in the single *asp* mutants (Figure 6F). Entire brain (D) and optic lobe (E) volume measurements (T-ratio) from WT, *asp* mutant, and *asp* mutants expressing UAS-P35 using the Insc-Gal4 and Elav-Gal4 drivers. For violin plots, solid red line represents the median, and the dashed lines denote interquartile range (IQR). Welch’s t-test. ns, P>0.05; *P≤0.05; ****P≤0.0001. Error bars represent standard deviation. Scale bars = 50 μm.

## DISCUSSION

Our primary motivation in this work was to examine the neurodevelopmental time course of MCPH and identify the genetic and cellular factors contributing to it, using the fly as a model system. Our neuronal cell counts coupled with accurate volume measurements of the CNS suggests that MCPH is already evident during the late neurogenic time window of development, which persists during later stages of neural development without significant changes to brain size and neuronal cell number. The key question is ultimately the fate of these cells—why are they not being made correctly? Work in vertebrates have identified defects in mitosis, cell cycle length control, centriole duplication, maintenance of apical complex proteins, and premature delamination of radial glial cells as contributors to *ASPM* MCPH [14,17,18,23], suggesting that the etiology of the disorder is complex and involves multiple genetic and cellular pathways operating in parallel.

The advantage of our transcriptome analysis is that it leverages the benefits of –omics-based approaches [19] and highlights many potential new pathways involved in *asp* MCPH, thus broadening the scope of the cellular basis of the disorder. However, untangling these multiple inputs is complicated in human MCPH patients, where access to viable tissue, especially during the neurogenic window of neurodevelopment, can be difficult to acquire [22]. One advantage of using the fly system is the ease at which living mutant tissue can be acquired at defined neurodevelopmental stages, and the availability of genetic tools to functionally assess candidate hits to identify the actual pathways and their relative contributions to the disorder.

The data presented here provides insights into this approach, both in terms of its advantages and its limitations. To our knowledge, this is the only bulk transcriptomic dataset available for *asp/aspm* microcephaly. A previous study used single cell RNA sequencing (scRNA-Seq) to examine the transcriptional profile of an *ASPM^-/-^* ferret model, although the data was used to determine relative proportions of neural progenitor cells rather than examine transcriptional differences between wildtype and mutant animals [23]. Thus, whether the transcriptional signatures we have identified here are conserved in vertebrates remains to be seen, although it provides a suitable dataset for comparison. However, one limitation is the lack of cellular resolution in our dataset. For example, even the neurogenic larval brain contains a significant amount of post-mitotic neurons and glial cells among its many neural stem cell types, limiting the ability to reveal cell-type specific transcription signatures [33, 49]. Such data would have allowed for deeper investigation into the cellular basis of the neurogenesis transcription factor network and immune system pathways, for example, and also help identify other contributing pathways that may have been filtered out in the bulk analysis.

Our co-expression-based module analysis was employed to partially overcome this limitation and provide a more thorough evaluation of the gene regulatory networks operating in *asp* mutant brains. We were encouraged by the fact that from a single GO-level perspective, the biological pathways identified in our function analysis (immune system, neurogenesis, proteolysis, stress & metabolic pathways, etc.) were also found in the module analysis. Also, the module analysis provided greater insight into how these disparate pathways might actually be coupled at the transcriptional level and integrated into multiple inputs operating in parallel.

However, it is important to note that the construction of these co-expressed networks are dependent on the input datasets used (microarray in this case) [34], and further refinement of them could be achieved through additional datasets, especially from the tissues of interest. Thus, there is a possibility that the modules containing a large number of genes (>200) and different GO associations could be further resolved into their own unique modules. Nonetheless, even though this would suggest an uncoupling at the expression level, a high enrichment of the individual modules as a whole would still support the conclusion of multiple biological pathways operating in parallel in *asp* mutant brains.

Another insight involves the genetic characterization. Our double and triple mutant analysis to evaluate the contribution of the immune system activation we consistently observed across neurodevelopment revealed a partial rescue of the brain size phenotype. We chose to focus on inflammation for reasons outlined above, including its involvement in other fly neurodegenerative phenotypes, and because work by Lemaitre and others have elucidated the molecular underpinnings of this response through mutational analysis in *Drosophila* and is thus fairly well characterized with a number of mutant alleles available [38–41,44,50,51]. Although our data suggests that inflammation can contribute to the *asp* MCPH size phenotype, when and how this occurs is currently not known. We did not perturb immune function in a developmental stage manner, thus it remains an open question as to whether its involvement is more critical during neurogenesis or the later time points. As for the how, factors such as cell number, size, shape, and packing density can affect overall brain volume in vertebrates [13, 52], although the extent to which inflammation can alter these properties is less defined. We tested apoptosis, a common outcome of immune system activation in the fly CNS [40,41,43,48] and a logical pathway for the reduction in total cell number and brain size that we observed in *asp* mutants, but found it surprisingly does not contribute to the *asp* MCPH phenotype either. It also remains an open question as to whether the NF-ΚB factors have other transcriptional targets unrelated to the immune response that could be responsible. This is especially true for *dif*, who has additional developmental patterning roles during embryogenesis [53] and whose brains are ∼10% larger than wildtype.

Another observation from this analysis is that the two prominent *asp* microcephaly phenotypes, tissue morphology (neuropil disorganization) and size, appear to be independent of each other. We currently do not know what is responsible for these morphology defects, although it may be a consequence of the disrupted larval neuroepithelium seen in *asp* mutant larva animals thought to be a result of disrupted actin cytoskeleton function [8]. Interestingly, we did find an enrichment of actin, myosin, and other muscle related factors in our network analysis for larval and adult stages, suggesting an underlying transcriptional effect on the actin cytoskeleton might also contribute to one or both of the MCPH phenotypes, although we have yet to test this directly. Nonetheless, our data suggest that tissue morphology and size can be uncoupled from one another, providing additional evidence that the etiology of MCPH is complex that multiple pathways affected by *asp* loss collectively contribute to brain growth and development.

One unexpected finding, but one that we speculate is the primary driver of the neurogenic defects and subsequent *asp* MCPH phenotype, was the global downregulation of a number of transcription factors that regulate key neurodevelopmental events, particularly neurogenesis and eye development. A number of studies from the Desplan lab and others have revealed an intricate transcription factor cascade consisting of spatially and temporally acting factors that ultimately control the transition from progenitor cells to a diverse suite of neurons in the optic lobe [33,49,54–57]. Our larval analysis highlighted many of these factors, including *eyeless (ey), Dichaete (D), tailless (tll), odd paired (opa), sloppy paired 1 (slp1), earmuff (Erm), runt (run), Optix,* and *twin of eyeless (toy)* [33]. We also found other transcription factors known to play important roles in eye, optic lobe, and lamina development including *sine oculus (so), rough (ro), glass (gl), pebbled (peb), spalt major (salm), anterior open (aop),* and *dachshund (dac)* [49,55,58–61]. Our *asp* mutants display retina, lamina, and optic lobe defects, the latter of which is the most significant contributor to the MCPH phenotype [16] and are consistent with mutant phenotypes reported for a number of these factors. *Eyeless* mutants, for example, show optic lobe morphological defects that are strikingly similar to those seen in *asp* mutants, including reduced medulla, lobula & lobula plate size and orientation, plus defects at the neuropil boundaries and ectopic fiber bundles [62]. Further work will be needed to determine the relative contribution of these spatial and temporal TFs in the *asp* MCPH phenotype and are being prioritized for follow-up.

In addition to these transcription factors, we also found an enrichment of Notch signaling related components that were also downregulated, particularly the *e(spl)* complex of transcriptional repressors [63]. Notch signaling’s role in neurogenesis is well-established, operating at multiple points along the temporal progression from neuroblasts to differentiated neurons to determine cell fate in both flies and vertebrates [12,26,64,65]. In the fly optic lobe, loss of function *Notch* and *Delta* mutants have a smaller medulla and lamina as a result of premature differentiation of neuroepithelial cells into medulla neuroblasts, with gain of function mutations showing an expansion of the neuroepithelial pool at the expense of neuroblasts [66]. Notch signaling has also been shown to directly regulate the expression of the temporal transcription factor *slp1* in the medulla [67], which was identified in our analysis. Notch also has a role in the terminal differentiation of neurons in the optic lobe of flies, where it cooperates with the temporal transcription factors mentioned above (e.g., *D, toy, run*) in a birth-order dependent fashion to generate neuronal diversity in the medulla [33,66,68]. Thus, disruption of Notch signaling in the brain could have a diverse range of cellular consequences that span the entire neurogenesis program of the developing optic lobe. Experiments are currently underway to address these questions, although it is also worth mentioning that our *asp* mutants phenocopy *Notch* mutants in the wing and bristles [69]. However, our module analysis also found many of the *e(spl)* genes coexpressed with genes of both known and unknown function, again suggesting that if Notch signaling contributes to the MCPH phenotype, it likely does so with multiple other inputs as well.

Lastly, the question of how mutations in *asp* could lead to a downregulation of a number of spatial and temporal optic lobe transcription factors and their regulatory networks remains unknown. One possibility is that Asp may have a direct role in regulating these genes and their networks by acting as a transcription or chromatin factor. Interestingly, Asp and its human ortholog (ASPM) have been shown to localize to the interphase nucleus of *Drosophila* meiotic I cells and human cultured cells [6,15,70], although whether it has a nuclear function (such as transcriptional regulation) remains to be explored. Given that the regulatory hierarchy of these temporal factors is complex and not fully understood [33], much more work will be needed to explore this. Another possibility is that Asp regulates the signaling pathways (e.g., Notch) that dictate proper spatial and temporal expression patterns of these factors through direct protein-protein interactions of pathway regulators, or perhaps even direct interaction with the tTFs themselves. More work will be needed to test these hypothesis and determine the relative contribution of these regulatory pathways and others in the etiology of *asp* MCPH.

## METHODS

### Fly stocks and husbandry

All stocks and crosses were maintained on standard cornmeal-agar media at 25°C. The *asp* mutant alleles (*asp^T25^* & *asp^Df^*) and *asp* transgenic rescue lines were previously described [5, 16]. The following lines were obtained from the Bloomington Stock Center: *w^1118^; Rel^E38^, e[s]* (BS#: 9458); *P{ry^+t7.2^=Dipt2.2-lacZ}1, P{w^+mC^=Drs-GFP.JM804}1, y^1^ w*; Dif^1^ cn^1^ bw^1^*(BS#: 36559); *w^1118^; P{w^+m*^=GAL4}repo/TM3, Sb^1^* (BS#: 7415); *w[*]; P{w[+mW.hs]=GawB}insc[Mz1407]* (BS#: 8751); *P{w[+mC]=GAL4-elav.L}2/CyO* (BS#: 8765); *y*[1] *w[*]; P{Act5C-GAL4-w}E1/CyO* (BS#: 25374); *w[*]; P{w[+mC]=UAS-p35.H}BH1* (BS#: 5072). The *UAS-H2Av::RFP* line was obtained from Laura Buttitta. Mutations were verified using Single-wing PCR [71] and Sanger Sequencing. All UAS- and Ubi-transgenic lines were generated in the *yw* mutant background by BestGene (Chino Hills, CA, USA).

### Developmental Timing & Brain Dissection

Animals were staged based on the following criteria: Wandering 3rd Instar Larvae were collected from non-crowded vials once they emerged from the food; P7 Pupae were collected as white prepupae, placed in a 35 mm dish with a moist Kim wipe, and allowed to develop for an additional 48 hours at 25°C; adult females were collected shortly after eclosion and aged for 3-5 days. Larval and pupal brains were rapidly dissected in fresh, room temperature SF900 media. Adults were dipped into 100% EtOH for 3-5 seconds prior to dissection in fresh SF900 in order to remove the waxy covering of the cuticle and allow for easier dissection. Dissected brains were maintained in fresh SF900 media until sample downstream processing for no longer than 15 minutes at room temperature. Four biological replicates consisting of 10 to 15 brains for each genotype and developmental stage were used for downstream processing.

### Total RNA extraction

Total RNA was extracted using Trizol® reagent (Invitrogen) following the manufacturer’s instructions. The RNA was resuspended in 12.5ul of sterile Milli-Q water and treated with Turbo^TM^ DNAse (Invitrogen) to eliminate DNA contamination. Quantity and quality of RNA were assessed using a Nanodrop 2000 spectrophotometer (Thermo Scientific, USA) and 1% (w/v) agarose gel electrophoresis, respectively.

### RNA Sequencing & Differential Gene Expression Analysis

Library preparation and RNA Sequencing was performed by Novagene (UC Davis, USA) using an Illumina NovaSeq 6000 platform. All samples had a RIN score ≥6.5. Sequencing generated a total of 881 million paired-end reads of 150 bp. Raw sequencing reads were trimmed and filtered using FastQC and Cutadapt with length cutoff of 20 nt [72, 73]. Surviving reads were mapped to the *Drosophila melanogaster* reference genome (dm6) using the RSubread package [74]. Gene expression levels were estimated using FeatureCounts software with default settings [75]. Differential gene expression (DGE) analysis was performed using the EdgeR Bioconductor package, filtering for CPM <1 [76]. Genes with a logF.C. ≥ 0.5/≤-0.5 and an adjusted P-value ≤0.05 were assigned as differentially expressed.

### Functional Annotation and Network Analysis

Identification of biologically relevant pathways and gene sets was performed using DAVID (v2021) and the EnrichmentMap & GeneMANIA plugins for Cytoscape (v3.9.1) [30–32,77,78]. Differentially expressed genes were first analyzed in DAVID using the Functional Annotation Tool. Parameters included 1) Functional Annotations: UP KW Biological Process, UP KW Cellular Component, UP KW Molecular Function; 2) Gene Ontology: GOTERM BP Direct, GOTERM CC Direct, GOTERM MF Direct; 3) Pathways: KEGG Pathway. Functional Annotation Clustering was then performed using default settings and a medium Classification Stringency. Functional Annotation Charts were also generated in DAVID using default settings, which served as the input for EnrichmentMap. Cutoff values using the Overlap Index test for each dataset included P-value: 0.05, FDR Q-value: 0.1, Overlap 0.6. Gene set nodes were then analyzed in GeneMANIA using default settings to visualize connectivity between genes based on previously published Co-expression, Predicted, Co-localization, Physical Interactions, and Genetic Interactions datasets.

### Independent Component analysis (ICA)

A second analysis pipeline was used to assess functional enrichment of differentially expressed genes (DEGs) based on *Drosophila*-specific gene expression signatures [34]. These gene expression signatures were identified through Independent Component Analysis (ICA) of 3,346 microarray datasets generated by the fly community and consist of 850 modules of co-regulated genes. ICA modules were encoded as a sparse non-negative 18,952 microarray probe sets by 850 modules weight matrix derived from a 18,952 microarray probe sets by 425 independent components ICA **S** matrix (see [34] for details). 850 sets of the most heavily-weighted co-regulated genes were created by discretizing (probe set weight ≥ 3) each of the ICA module matrix vectors and used to determine module GO term and KEGG pathway enrichments (Supplementary File 4). To project the expression data in this report onto the ICA module matrix, the matrix probe set features were first converted to FBgn features compatible with EdgeR logF.C. vectors using the biomaRt package. ICA module matrix weights for the relatively few probe sets that mapped to the same FBgn were averaged to yield the 13801 FBgn by 850 modules ICA module matrix used here. Prior to projection onto the FBgn-based ICA module matrix, EdgeR outputted genotype pair FBgn logF.C.s vectors were mean 0 centered and variance 1 scaled. After restricting the ICA module matrix and logF.C. vectors to the FBgn features shared between them, scalar projection of the logF.C. vectors onto the ICA module matrix was performed as described previously [34]. In addition, one hundred random permutations of each genotype pair logF.C. vector (100 shuffles of FBgn label to logF.C. value relationships) were also projected onto the ICA module matrix. The mean and standard deviation of random permutation scalars per module were used to compute enrichment Z-score and P-value significance values for each module in a genotype pair logF.C. vector. A module was considered significantly enriched if it obtained a Z-score ≥+3/≤-3 and a P-value≤0.05.

### Gene expression: RT-qPCR

cDNA was synthesized via the iScript™ cDNA Synthesis Kit (BioRad, USA) using 1 μg of total RNA. The resultant cDNA was then used 1:10 in the qPCR reaction, which consisted of iQ™ SYBR® Green Supermix (BioRad) and gene-specific primers. Primers are available upon request. The following cycling parameters were used: 95 °C for 5 min, 40 cycles of 95 °C for 10s, and 60 °C for 45s ending with a melting curve. Relative gene expression was analyzed by the multiple reference gene method. Elongation factor 1-alpha (ef1-α) and RP49 were used as the internal reference genes. Relative quantities and data normalization followed the formulas detailed in Hellemans et al. [79]. The comparative Cq (ΔΔCq) method was employed to calculate the relative expression ratios (RER). Three technical replicates were performed for each of the four independent biological replicates assayed. Statistical analysis was performed using ANOVA, Bonferroni’s post test, and t-test when applicable using PRISM 9 (GraphPad Software, USA).

### Immunohistochemistry & Antibodies

Larval and adult brains were carefully dissected in SF900 media, transferred to 1.5 mL tubes containing 1.2% paraformaldehyde (PFA) diluted in 1xPBS, and fixed for 24 hours while nutating at 4°C. Fixed brains were then washed in 0.5% PBT for 3×15 min at RT, blocked for 1.5 hours in 0.5% PBT with 5% normal goat serum, and nutated with anti-Bruchpilot antibody (1:30, nc82, DSHB), anti-cleaved Death Caspase-1 (DCP-1, 1:100, Cell Signaling Technology), anti-β-Tubulin (1:250, E7, DSHB), or anti-phospho-Histone H3 (Ser10) (1:1000, Millipore Sigma) for 4 hours at RT before transferring to 4°C for two overnights. Brains were then washed using 0.5% PBT (4×15 minutes) and incubated 1:500 with secondary antibody (anti-mouse/rabbit Alexa-488/647, Invitrogen/ThermoFisher) for 4 hours at RT before transferring to 4°C for three overnights. Brains were again washed using 0.5% PBT (4×15 min) and stained with DAPI (0.1 ug/mL). Following a post-fixation step with 4% PFA at RT for 4 hrs, the brains were washed using 0.5% PBT (4×15 min) and mounted on poly-L-lysine coverslips. The coverslips were transferred through an ethanol dehydration series (30%, 50%, 75%, 95%, 100% x3), cleared using xylene, and mounted in DPX mounting media. Slides were allowed to cure for at least 2 overnights before imaging.

### Microscopy

All imaging was conducted on an Olympus IX83 microscope fitted with a Yokogawa CSU-W1 Dual Disk SoRa, dual Hamamatsu Orca Flash 4.0 V3 sCMOS cameras, and Plan S-Apo 40x & 100x Si Oil Objectives (NA 1.25, 1.35) operated by cellSens software.

### Apoptotic Cell Quantification

DCP-1 positive cell counts were obtained using the surface counter feature in Imaris (v9.9.1). Uniform threshold values were established for the intensity of cells positive for DCP-1 based off positive control and negative control brains. Other parameters were established prior to analysis including: cell seed diameter for completing a morphological split of touching or grouped cells, surface smoothing, quality evaluation of the detected surfaces, and the minimum number of voxels per object. All raw cell counts were further normalized by dividing the cell count for the optic lobe by the total volume of the optic lobe which was calculated using the segmentation feature in Dragonfly software (Object Research Systems, v2020.2.0.941). Thresholding was utilized during segmentation in Dragonfly to ensure measurements only included the optic lobe of interest.

### Microcomputed Tomography (u-CT)

Sample preparation, imaging, and analysis were carried out as previously described [16, 80]. I2KI was used as the contrast agent. Samples were scanned using a SkyScan 1172 desktop scanner controlled by Skyscan software (Bruker). X-Ray source voltage and current settings: 40 kV, 250 μA, 4 W of power. A cooled 14-bit 11Mp (402×2688) CCD detector coupled to a scintillator was used to collect X-rays converted to photons. Medium camera settings at an image pixel size of 2.95 μm were used for fast scans (∼20 min), which consisted of about 300 projection images. Frame averaging used was 2. Tomograms were generated using NRecon software (Bruker MicroCT, v1.7.0.4). Dragonfly (Object Research Systems, v2020.2.0.941) was used for manual brain segmentation and determination of brain and optic lobe volume.

### Flow Cytometry

We followed the method of Nandakumar et al. to obtain total cell counts from adult brains [29]. Single adult brains were dissected in SF900 media and transferred to a sample tube containing 90 μl Trypsin-EDTA, 10 μl 10X PBS, and 2 μl of Vybrant™ DyeCycle™ Violet stain (ThermoFisher Scientific). Brains were incubated for 20 minutes at room temperature before being triturated using a low retention P200 pipette tip for 90 seconds. 400 μl of the Trypsin-EDTA-DyeCycle solution was then added to allow for additional digestion of the tissue for 45 minutes at room temperature. The sample was then diluted with 500 μl of 1x PBS and gently vortex to resuspend the cells. Samples were run at a slow flow rate (∼280 events/sec or 1ml per 15 minutes) using a BD FACSMelody™ (BD Biosciences) until the entire volume of the tube was empty. Data was analyzed using FlowJo software (v10.7.2, BD Biosciences). Events were gated using relative size (FSC) and granularity (SSC). To establish fluorescent vs. non-fluorescent populations of events, unlabeled Oregon-R brains were used to establish the maximum non-fluorescence threshold based on intensity. Events that populated above this threshold in the DyeCycle labeled samples were gated as positive and counted. We verified that the majority of these events consisted of brain cells by subsequent flow sorting and immunofluorescence microscopy using an anti-lamin antibody (1:100, ADL101, DSHB) to visualize the nuclear envelope. We also verified this population using an UAS-H2Av::RFP line driven by Actin-Gal4, which were double positive for both DyeCycle Violet and RFP signal indicating intact fluorescent nuclei (Supplementary Figure 2).

## ACKNOWLEDGEMENTS

We thank Laura Buttitta for sharing fly strains, advice, and protocols for flow cytometry of fly brains, Jason Gigley for training and support on the FACSMelody™, and Missy Stuart for maintaining fly stocks and S2 cells. This work was supported by grants from the National Heart, Lung, and Blood Institute of the National Institutes Health (1K22HL137902-01) and an Institutional Development Award (IDeA) from the National Institute of General Medical Sciences of the National Institutes of Health under Grant #2P20GM103432. SC and SF are supported by the USDA National Institute of Food and Agriculture, Hatch project #1012152. MBC is supported by the Wyoming Research Scholars Program. Any opinions, findings, conclusions, or recommendations expressed in this publication are those of the author(s) and do not necessarily reflect the view of the NIH, National Institute of Food and Agriculture (NIFA) or the United States Department of Agriculture (USDA).

## COMPETING INTERESTS

The authors declare no competing interests.

## DATA AVAILABILITY STATEMENT

All raw and processed sequencing data generated in this study will be submitted to the NCBI Gene Expression Omnibus (GEO; https://www.ncbi.nlm.nih.gov/geo/) prior to publication, at which time an accession number will be made available.

---

**Supplementary Figure 1.**
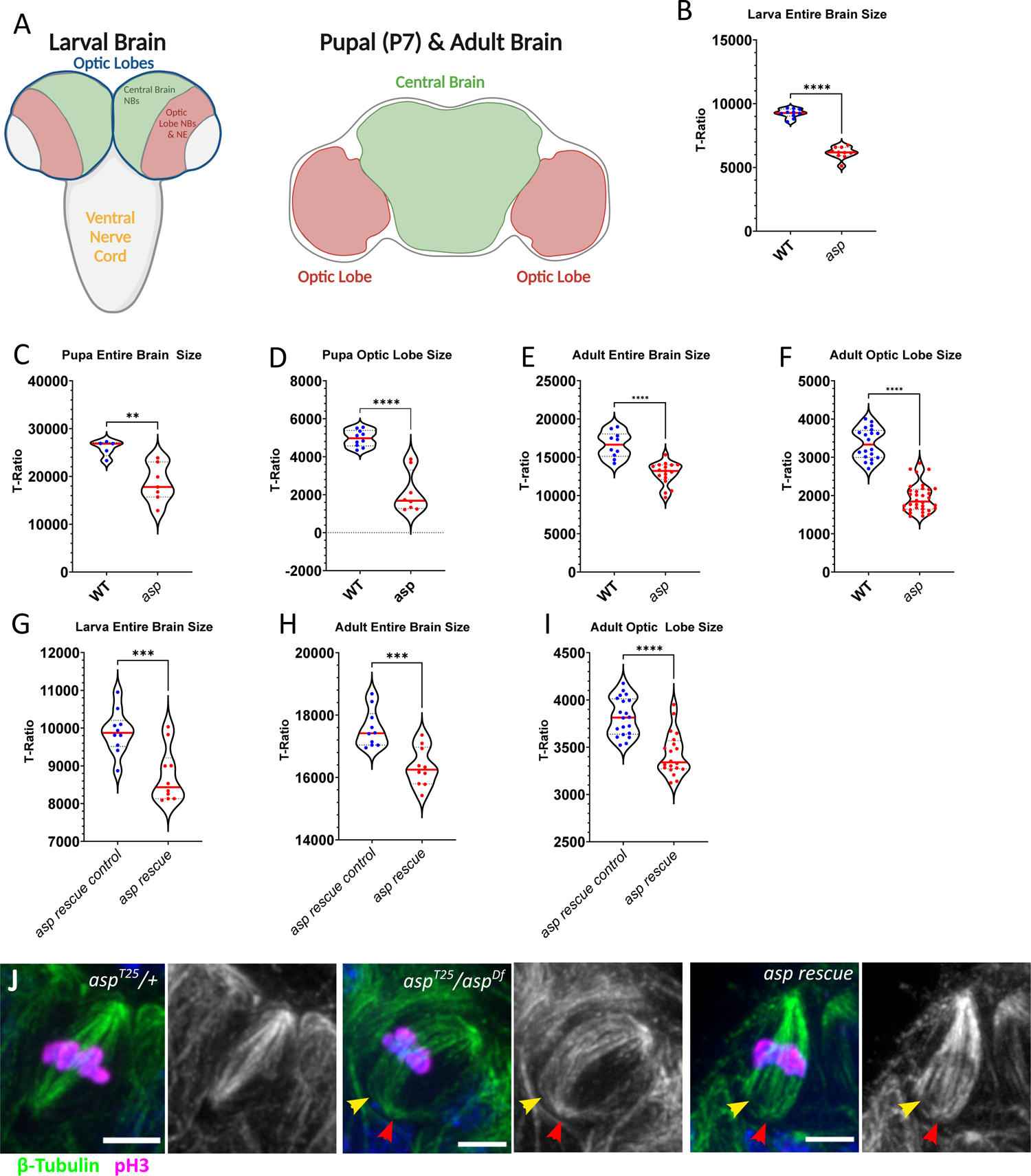
Related to Figure 1. Brain size measurements and mitotic spindle morphology. (A) Cartoon of the larval and pupal/adult brain. The optic lobes of the larval brain consists of two neuroblast (NBs) regions that will give rise to distinct neuron and glial populations in the adult brain. Central brain NBs (green) make neurons and glia for the central brain of the adult, optic lobe NBs (also known as medulla neuroblasts) and neuroepithelial cells (red) generate the neurons and glia of the adult optic lobe. For μ-CT measurements, the ‘entire brain’ consists of both larval optic lobes (blue outline) segmented as a whole, while the pupal and adult entire brain consists of both optic lobe regions plus the central brain. ‘Optic lobe’ measurements were taken from individually segmented optic lobes of the pupa and adult. The larval ventral nerve cord was not included in the size analysis, although it was included in the flow cytometry analysis due to the difficulty in accurately separating it from the optic lobes during dissection. μ-CT measurements of wildtype (WT, *asp^T25^/+*) and *asp* mutant (*asp^T25^/asp^Df^*) volume from (B) larva entire brain, (C) pupa entire brain, (D) pupa optic lobe, (E) adult entire brain, (F) adult optic lobe. μ-CT measurements of *asp rescue control* (*ubi-GFP::asp^MF^/+; asp^T25^/+*) and *asp rescue* (*ubi-GFP::asp^MF^/+; asp^T25^/asp^Df^*) volume from (G) larval entire brain, (H) adult entire brain, and (I) adult optic lobes. Data is represented as the T-ratio (brain volume normalized to overall body size) and each dot represents a single brain. These values were used to generate the ‘normalized’ graphs in Figure 1B, 1C. For violin plots, solid red line represents the median, and the dashed lines denote interquartile range (IQR). (J) Metaphase mitotic spindles of larval neuroepithelial cells undergoing symmetric division, labeled with anti-β-tubulin to visualize microtubules and anti-pH3 to visualize mitotic chromosomes in WT (*asp^T25^/+*), *asp* mutant (*asp^T25^/asp^Df^*), and *asp rescue* (*ubi-GFP::asp^MF^/+; asp^T25^/asp^Df^*) brains. Yellow arrowhead denotes unfocused spindle poles, red arrowhead indicates partially detached centrosomes. n≥5 brains; Welch’s t-test. ns, P>0.05; **P≤0.01; ***P≤0.001; ****P≤0.0001.

**Supplementary Figure 2.**
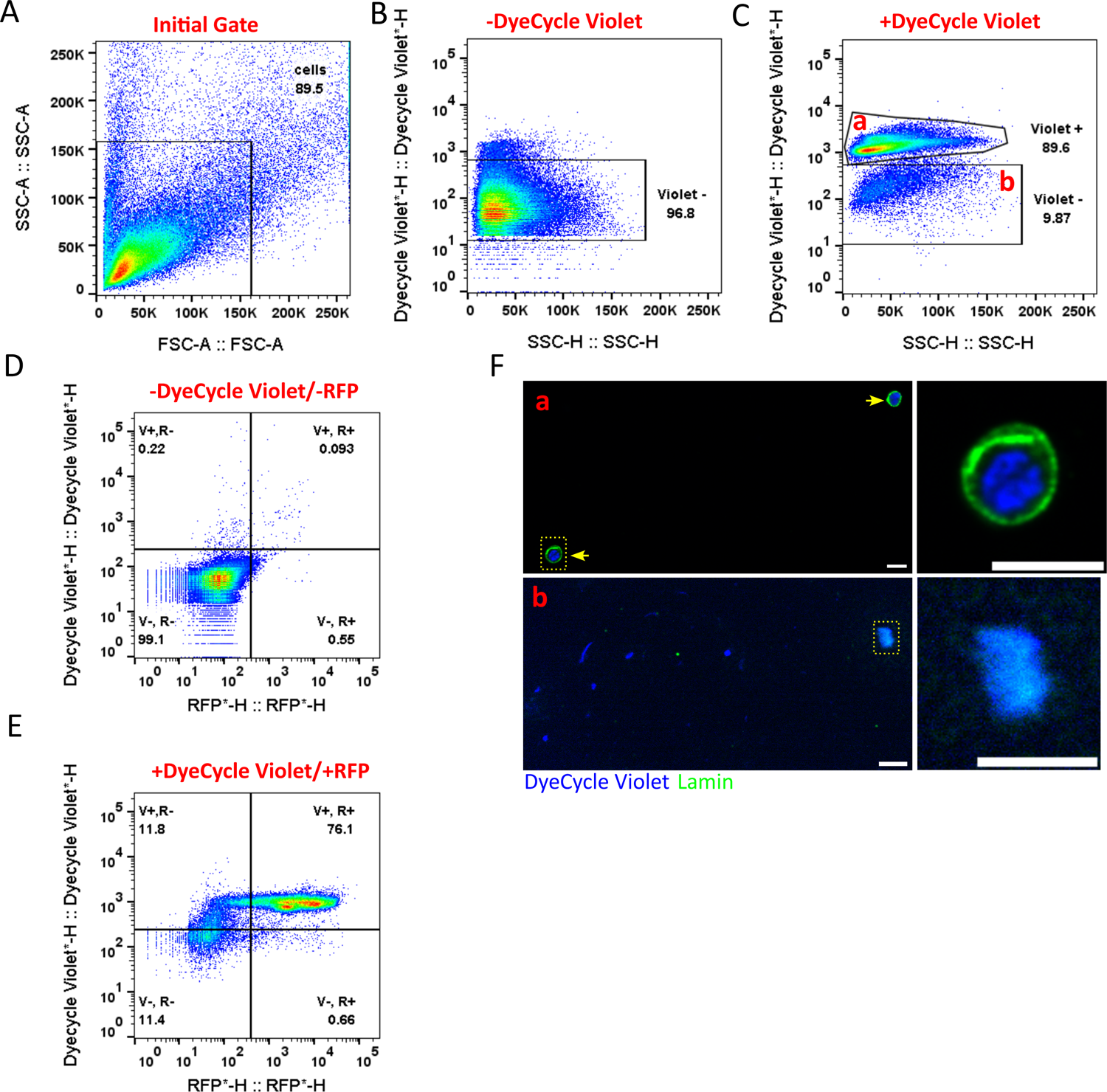
, Related to Figure 1. Validation of flow cytometry for neuronal cell counts from single brains. (A) Forward scatter (FSC-A) vs. Side scatter (SSC-A) plot from a single dissociated adult brain. Boxed area indicates the initial gate of events used for subsequent analysis. (B) DyeCycle Violet™ vs SSC-H of the gated area in (A) from an unlabeled (-DyeCycle Violet) brain, which serves as a negative control to establish the fluorescence threshold of a positive event (labeled nuclei). (C) DyeCycle Violet™ vs SSC-H of a labeled (+DyeCycle Violet) brain. The gate labeled ‘a’ represent intact nuclei, as judged by FACS sorting and subsequent immunostaining of this population using an anti-lamin antibody (Panel F, top panels) and was used to determine the number of cells (events) shown in Figure 1E-H. The gate labeled ‘b’ represents debris and was not included in the analysis (Panel F, bottom panel). A second validation was performed using an Actin-Gal4/UAS-H2Av::RFP strain that was either not co-labeled (D) or was co-labeled with DyeCycle Violet™ (E). Events displaying a high intensity in both the RFP and DyeCycle violet channels (V+, R+) were intact nuclei, and correspond to the ‘a’ gate shown in (C). (F) Microscopy images of the events sorted from the ‘a’ and ‘b’ gates in panel (C). Yellow arrowheads denote intact nuclei as judged by intact nuclear lamin staining, which was not observed in the ‘b’ population. Scale bars= 5 μm.

**Supplementary Figure 3.**
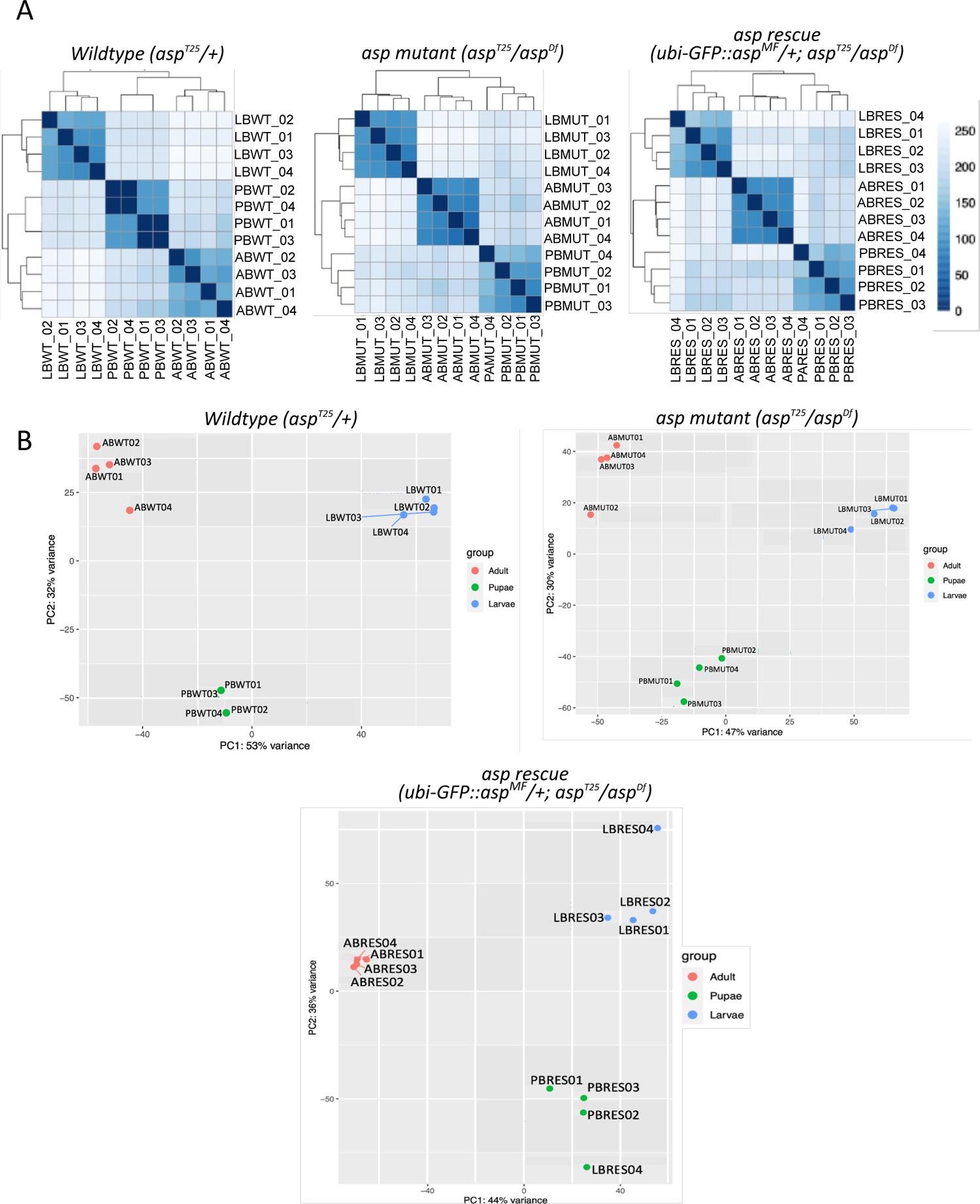
Related to Figure 2. Sample variability between RNA-Seq datasets. (A) Sample-to-Sample distances for each genotype (Wildtype, *asp* mutant and *asp* rescue) and developmental stage, calculated using edgeR. (B) PCA analysis of each genotype and developmental stage. PC1 & PC2 describe the majority (>75%) of the variability in the datasets.

**Supplementary Figure 4.**
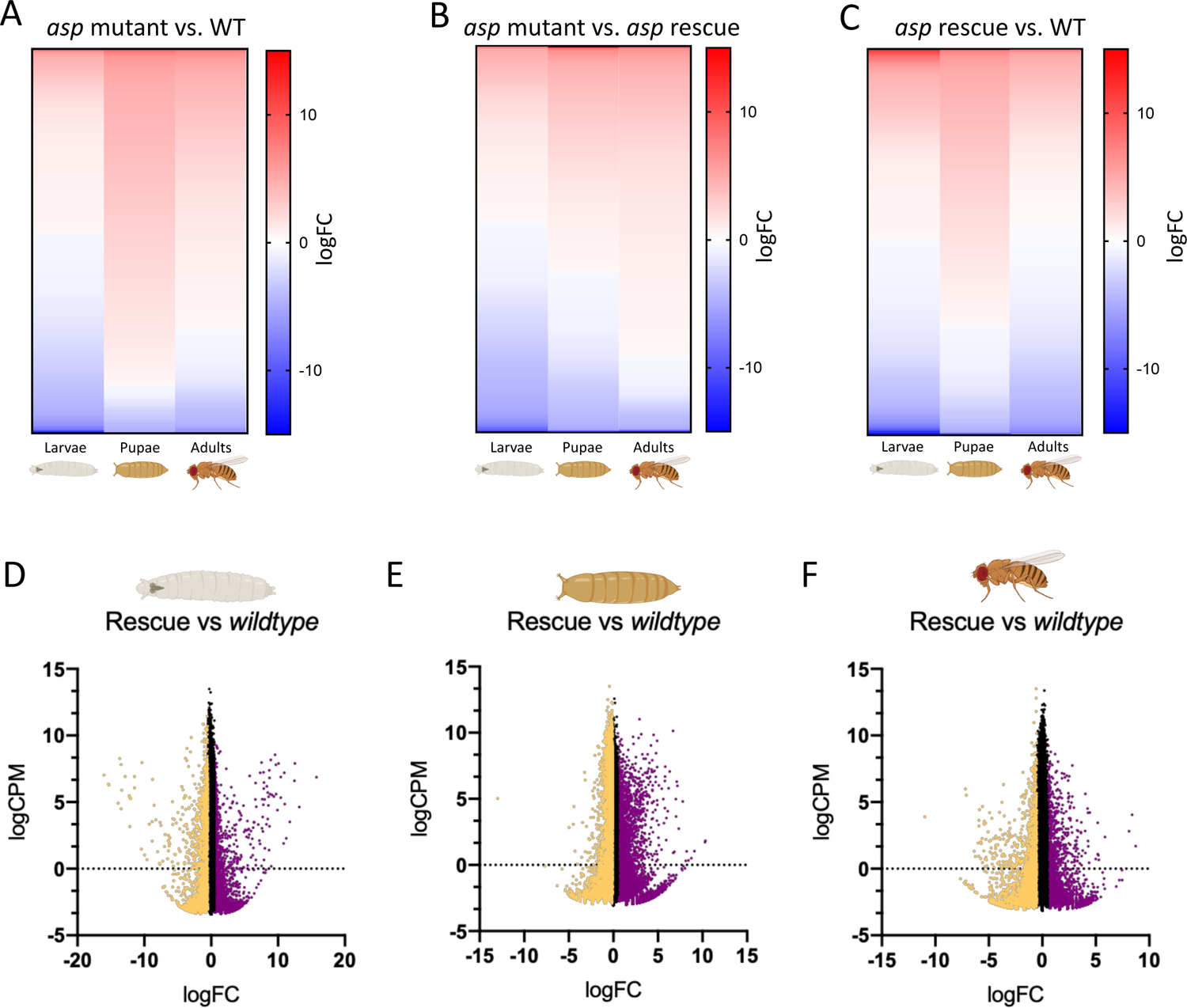
Related to Figure 2. Transcription trends across development. Heatmap representation of all DES across all developmental stages (larvae, pupae, adults) for the (A) *asp* mutant vs. WT, (B) *asp* mutant vs. *asp* rescue, and (C) *asp* rescue vs. WT comparisons, colored by logF.C. (red=>.5, blue =<-.5). logCPM vs logF.C. plots for the *asp* rescue vs. WT comparisons from (D) larvae, (E) pupae, and (F) adult brains. WT=wildtype.

**Supplementary Figure 5.**
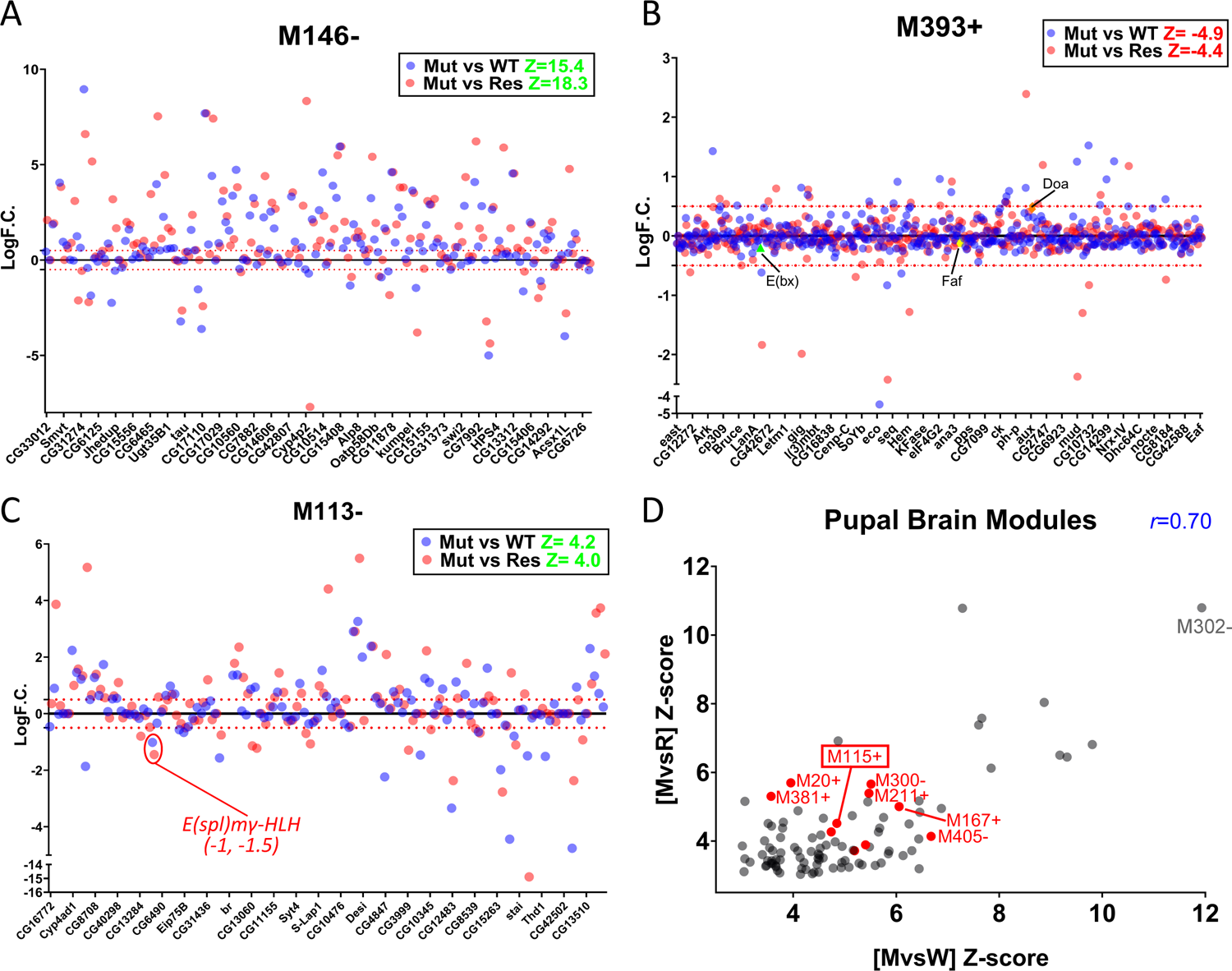
, Related to Figure 5. ICA module analysis to identify co-expressed gene signatures in *asp* mutant brains. Plot of logF.C. vs. gene for the (A) larval M_146-,_ (B) adult M_393+_, and (C) larval M_113-_ modules. Z-score enrichment is shown in green font for each genotype comparison. Each dot is colored based on the mutant vs wildtype (Mut vs WT, blue) and mutant vs rescue (Mut vs Res, red) value. Red dotted line indicates the 0.5/-0.5 logF.C. position. Not all gene names are included on the x-axis for clarity. Relevant genes (*doa, E(bx), faf, e(spl)mγ-HLH*) are highlighted. (D) Scatterplot showing significantly enriched co-expression modules identified through ICA analysis of the [M vs W] and [M vs R] DGE datasets from pupal brains. Each module is plotted as a point based on its Z-score enrichment from each analysis. Only significant modules with a Z-score of >3/<-3 are shown. Co-expressed modules containing at least one immune system-related GO term are colored red, with M_115+_ outlined with a red box. A subset of the more significantly enriched (Z-score ≥10) modules are also labeled in gray font. The Pearson’s Correlation Coefficient (*r)* is shown in blue font (two-tailed P<0.0001).

**Supplementary Figure 6.**
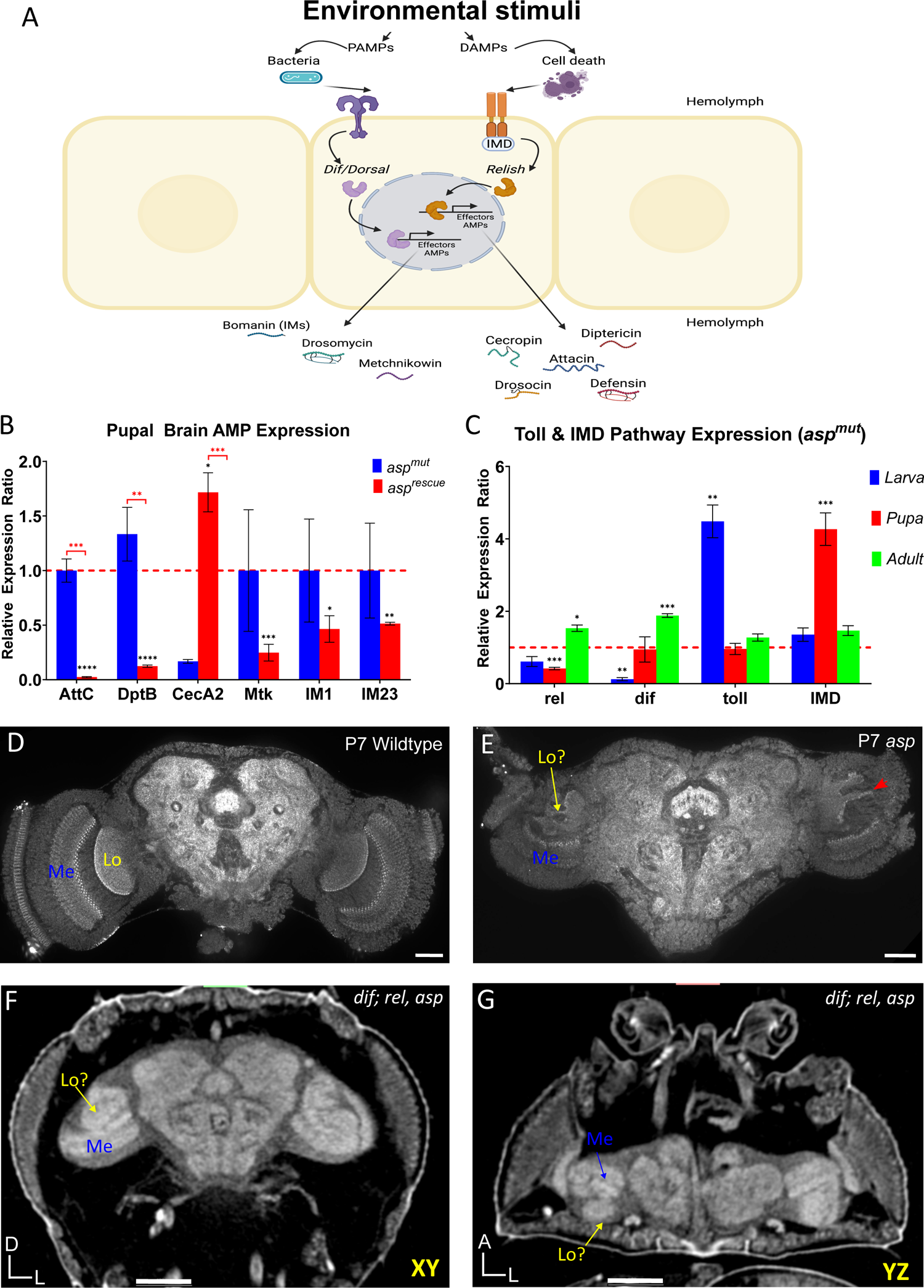
Related to Figure 6. Characterization of the fly CNS immune response. (A) Cartoon diagram of the insect immune response. Toll and IMD receptors respond to distinct stimuli (classified as PAMPs and DAMPs) and rely on the NF-ΚB factors Relish, Dorsal, and Dif to activate downstream effectors of the antimicrobial family of peptides (Bomanins, Attacins, etc). (B) AMP expression (qPCR) during pupal stages in *asp* mutant and *asp* rescue animals. Data is shown as the relative expression ratio (RER), red dotted line indicates the RER value for wildtype, normalized to 1. Expression in *asp* mutant brains is shown in blue, *asp* rescue in red. (C) qPCR analysis of the upstream Toll and IMD pathway components throughout development in *asp* mutants from larva (blue), pupa (red), and adults (green). Data is shown as the relative expression ratio (RER), red dotted line indicates the RER value for wildtype, normalized to 1. Confocal imaging of P7 pupal brains labelled with α-brp (nc82) to visualize the neuropil in (D) wildtype (*asp^T25^/+*) and (E) *asp* mutants (*asp^T25^/asp^Df^*). Red arrowheads point out disrupted neuropil boundaries. High Resolution μ-CT tomograms of *dif^1^; rel^E38^, asp* triple mutant showing an (F) XY plane and (G) YZ plane highlighting severe optic lobe neuropil disorganization identical to the *asp* mutant [16] (not shown). Multiple attempts were made to image triple mutant brains using confocal microscopy, but the ‘stickiness’ of these brains caused aggregation in the tubes and prevented proper mounting. Medulla (Me) and Lobula (Lo) neuropil regions are labeled. Body axes in each μ-CT view denoted as D, dorsal; L, left; A, anterior. Welch’s t-test. ns, P>0.05; **P≤0.01; ***P≤0.001; ****P≤0.0001. Error bars represent standard deviation. Scale bars = 50 μm (D, E) and 100 μm (F, G).

**Supplementary Figure 7.**
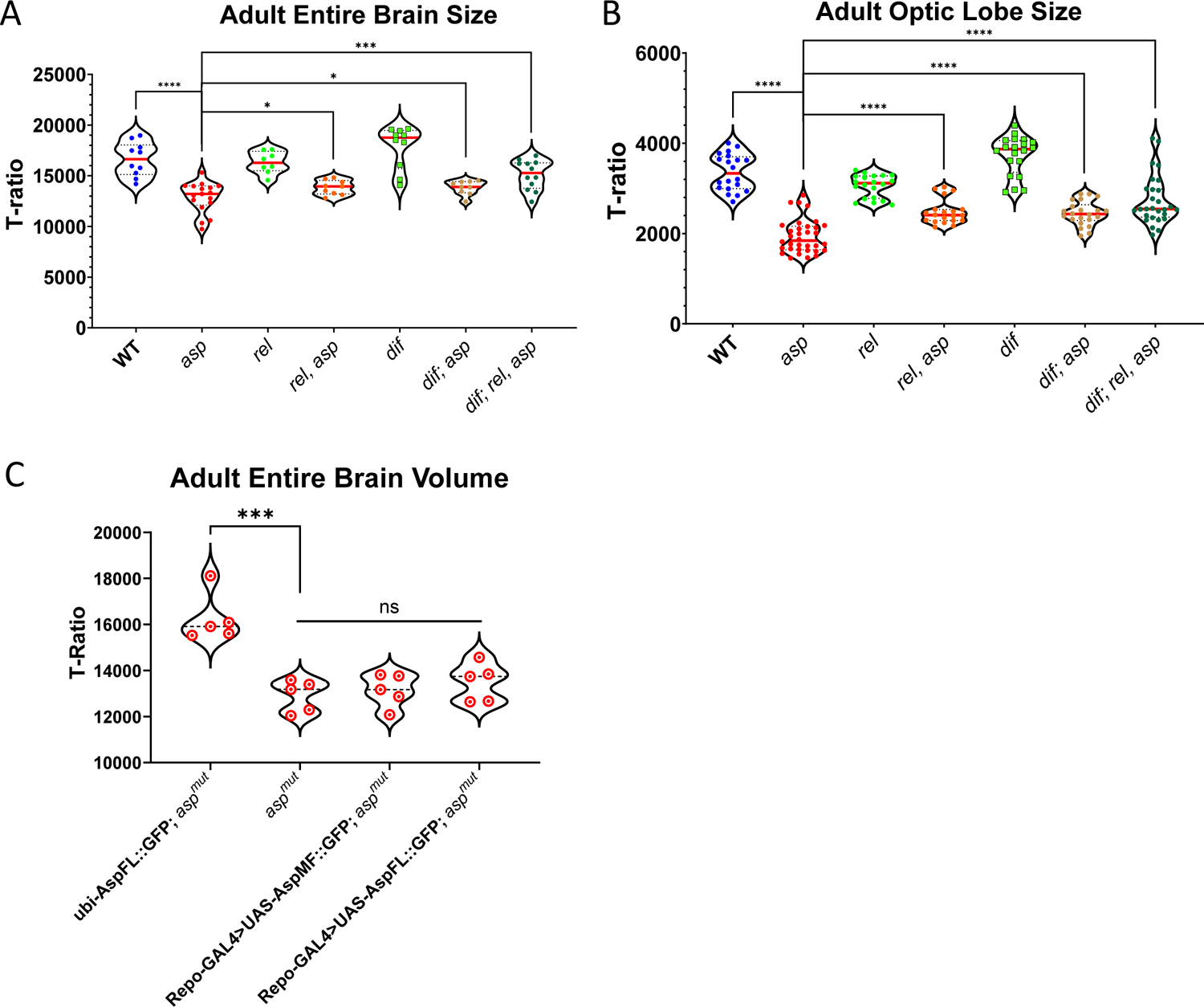
Related to Figure 7. Brain size measurements derived from μ-CT imaging. Individual adult (A) entire CNS volume and (B) optic lobe values (T-Ratio) used to generate the bar graphs in Figure 7 from the double and triple mutant genetic analysis. (C) Genetic rescue experiment for the entire adult brain using Repo-Gal4 to drive UAS-N-terminally tagged GFP Asp^MF^ and Asp^FL^ transgenes in glial cells in the *asp* mutant background. Ubi-Asp^FL^, driven constitutively under control of the ubiquitin promoter, serves as the “rescue” control. For violin plots, solid red line represents the median, and the dashed lines denote interquartile range (IQR). n≥5 brains; Welch’s t-test. ns, P>0.05; **P≤0.01; ***P≤0.001; ****P≤0.0001.

## Notes

### Competing Interest Statement

The authors have declared no competing interest.

